# A network of syringe-injectable bioelectronics for closed-loop therapeutic rescue

**DOI:** 10.64898/2026.11.10.765435

**Authors:** Joy Jackson, Wonbum Sohn, Lolita Testu, Ramy Ghanim, Maryam Aamir, Maya Angia, Alaz Cig, Sean Healy, Zachary Lee, Jason Y Chin, Julia Z Ding, Jihoon Park, Jannatim Maisha, Yoon Jae Lee, Alex Abramson

## Abstract

An on-body network of sensors and drug delivery implants could transform treatments for rapid-onset morbidities that require bystander intervention. During opioid overdoses, for example, lone and incapacitated victims often cannot self-administer life-saving naloxone in the minutes before respiratory collapse. However, the bulkiness of existing closed-loop systems threatens patient acceptance. Here, we present the Rapid Emergency Subcutaneous platform (RES-Q), a network of wearable sensors and syringe-implantable autoinjectors for closed-loop therapy. RES-Q implants are similar in size to FDA-approved drug rod implants and fit inside 6-gauge needles to facilitate outpatient administration. Informed by implant fibrosis fracture mechanics, RES-Q employs miniature jet injectors that deliver 0.4 mg human-relevant doses of naloxone through fibrotic encapsulation for rapid uptake. RES-Q implants communicate with wearable sensors through tissue conductivity, resulting in 25 nA standby current draws that ensure long-term functionality. In swine, we sequentially trigger multiple implants using ionic tissue communication. In rats, RES-Q detects fentanyl overdoses using wearable photoplethysmography sensors and performs autonomous, dose-titratable rescue in <6 minutes.

## Introduction

The opioid epidemic constitutes a widespread public health crisis in the United States, with more than 800,000 overdose deaths since the late 1990s^1^. Distribution programs for intranasal naloxone, a rescue drug that reverses overdose symptoms, are a primary harm reduction strategy. However, debilitating symptoms often prevent self-administration, and studies show 75% of deaths occur when an individual is alone^2^. Even with bystanders present, more than 50% fail to recognize the signs of a fatal overdose^3^. Naloxone administration within minutes can significantly mitigate the risk of brain injury or death^1, 4^; thus, new harm reduction strategies are critical.

In a recent survey of individuals who use opioids, more than 75% expressed interest in a long-term, closed-loop rescue device in the case of relapse^5^. However, 89% had concerns about device size and preferred the smallest device possible^5^. Current long-acting therapeutics for opioid use disorder include injectable drug depots and implantable drug rods measuring up to 2.5 x 26 mm^6^. Patients describe the discrete nature of these treatments as a benefit^7^, suggesting potential acceptance for a closed-loop rescue device of similar size. Many treatment centers are already staffed with the resources necessary for administering long-acting injectables, supporting the integration of an additional syringe-injectable device into existing clinical workflows. A syringe-injectable, closed-loop drug delivery system could also act as a platform technology for other emergency therapeutics, including glucagon delivery for patients suffering from severe hypoglycemia^8^ or epinephrine delivery for patients suffering from an allergic reaction^9^.

Existing experimental closed-loop devices for overdose rescue^10, 11^ and triggerable release^12–16^ necessitate invasive surgical implantation due to size and energy constraints associated with sensing, communication, and actuation that limit clinical translation. Communication protocols such as Bluetooth and Near Field Communication, for example, require large on-board antennas and microcontrollers. Furthermore, maintaining active communication using these methods reduces battery lifetimes to days or weeks^17^. Yet, without wireless communication, sensors and actuators must be directly wired together, increasing size. Overcoming fibrotic encapsulation, a form of scar tissue which impedes drug diffusion around implanted devices, can also involve bulky actuation components^18,19^. If unaddressed, fibrosis has been shown to slow subcutaneous naloxone uptake for the first 20 minutes after delivery, resulting in insufficient plasma drug concentrations for immediate overdose rescue^10^. Because the power required to break through the capsular tissue is not well characterized, scientists regularly engineer overpowered and oversized centimeter-scale pumps or motors to mechanically disturb or puncture fibrotic tissue during drug delivery^14, 20–22^. Other techniques for controlling drug delivery, including living cellular implants^23–25^, responsive hydrogels^26, 27^, and engineered microparticles^28^, also try to resolve the challenge of controlled burst release. However, they suffer from additional constraints including premature leakage, inferior burst release kinetics, and/or a lack of sensor-driven control (Table S1).

Here, we introduce the Rapid Emergency Subcutaneous (RES-Q) platform, a closed-loop network with syringe-implantable autoinjectors, that can detect and rescue an opioid overdose in less than six minutes *in vivo*. We harness the ionic conductivity of the human body^29^ to transmit signals directly through tissue from wearable overdose detection sensors to implants that hold 16 µL of a clinically-relevant 0.4 mg naloxone payload. Upon overdose detection, RES-Q implants deliver life-saving, spring-powered jet injections capable of cutting through fibrotic encapsulation to ensure long-term functionality (Fig. 1a-f). Compared to other experimental devices that demonstrate overdose rescue, RES-Q implants are 25-64x smaller, fit fully within 6-gauge needles, and improve theoretical battery lifetimes from weeks to years by only requiring 25 nA standby current draws (Table S2)^10, 11^. The syringe-implantable form factor and power efficiency result from miniaturized tissue-mediated ionic communication and electrolysis-pump-based triggering mechanisms, with deployment and confirmed functionality *in vivo* in swine and rodent models. To engineer a jet injection with sufficient power to break through fibrotic encapsulation without overshooting the actuator’s required power and size, we mechanically characterized, modeled, and performed a sensitivity analysis on the fracture mechanics of a jet injection through explanted fibrotic tissue (Fig. 1g). In rodent opioid overdose models, our system enables sequential, dual-stage delivery that controls naloxone dosage amounts, provides comparable pharmacokinetics to manual subcutaneous injections, and enables autonomous rapid overdose reversal in under six minutes.

**Fig. 1:**
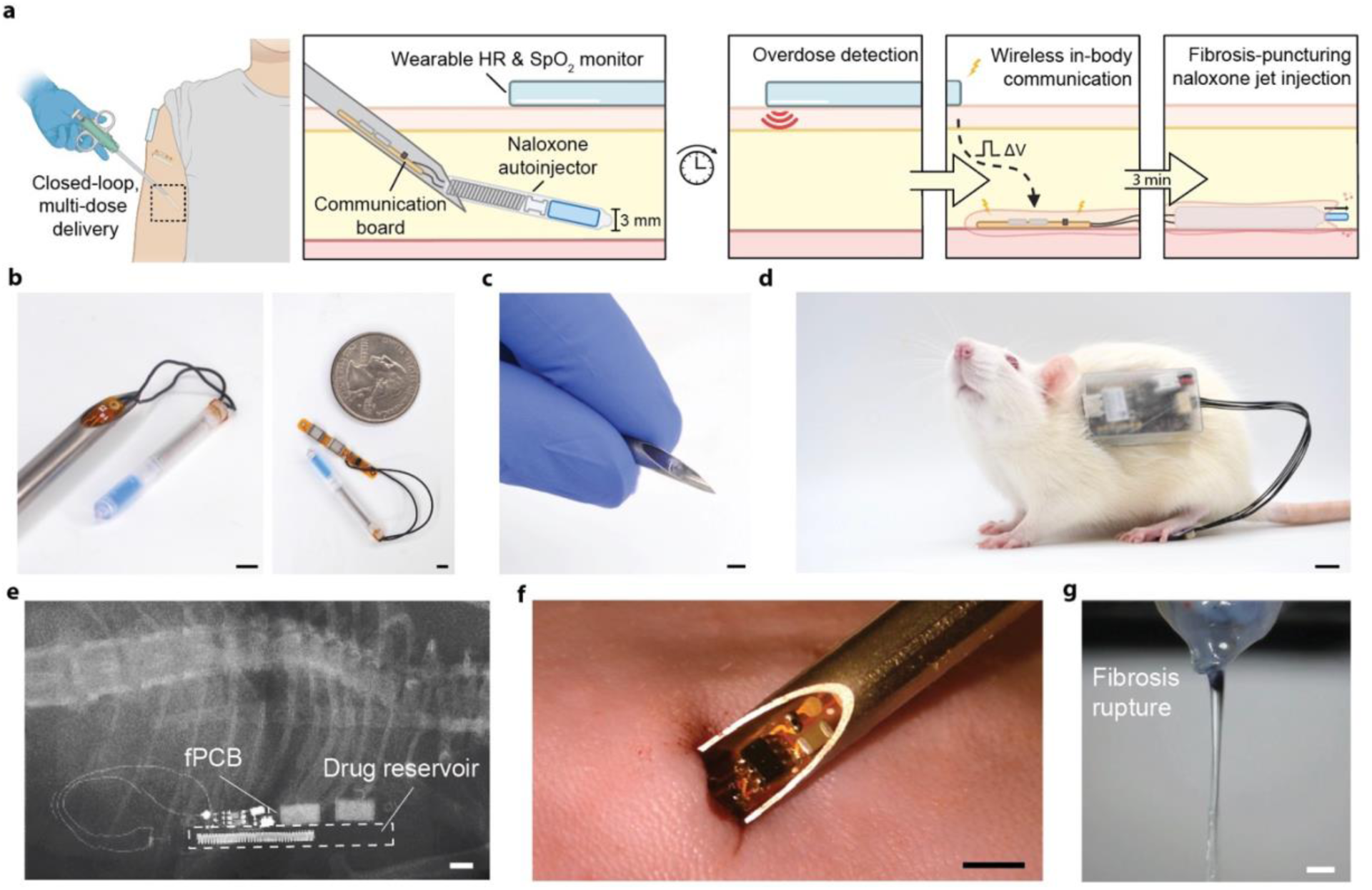
The Responsive Emergency Subcutaneous (RES-Q) network for closed-loop emergency drug delivery. **a**, System overview. HR = Heart rate. SpO_2_ = Peripheral capillary oxygen saturation. V = Voltage. **b-c**, Syringe-injectable drug delivery device (scale bar, 3 mm). **d,** Wearable vitals monitor on a rat (scale bar, 10 mm). **e**, Computed tomography (CT) scan of the drug delivery device implanted in a rat (scale bar, 3 mm). fPCB = Flexible printed circuit board. **f**, Syringe-implantation of the drug delivery device into a rat (scale bar, 3 mm). **g,** Rupture of fibrotic tissue via jet injection from the RES-Q device (scale bar, 1 mm). This image is also shown in panel 2f alongside jet injection simulations. Illustrations in **a** created with BioRender.com.

## Results

### Experimental and computational characterization of jet injection through fibrosis

To minimize the size of our syringe-implantable actuator, we found the minimum velocity necessary for a drug delivery jet to penetrate through fibrotic tissue that encapsulates long-term subcutaneous implants (Fig. 2a). We achieved this by building a computational fibrosis jet injection model informed by *ex vivo* mechanical characterization experiments on harvested fibrotic tissue (Fig. 2b).

**Fig. 2:**
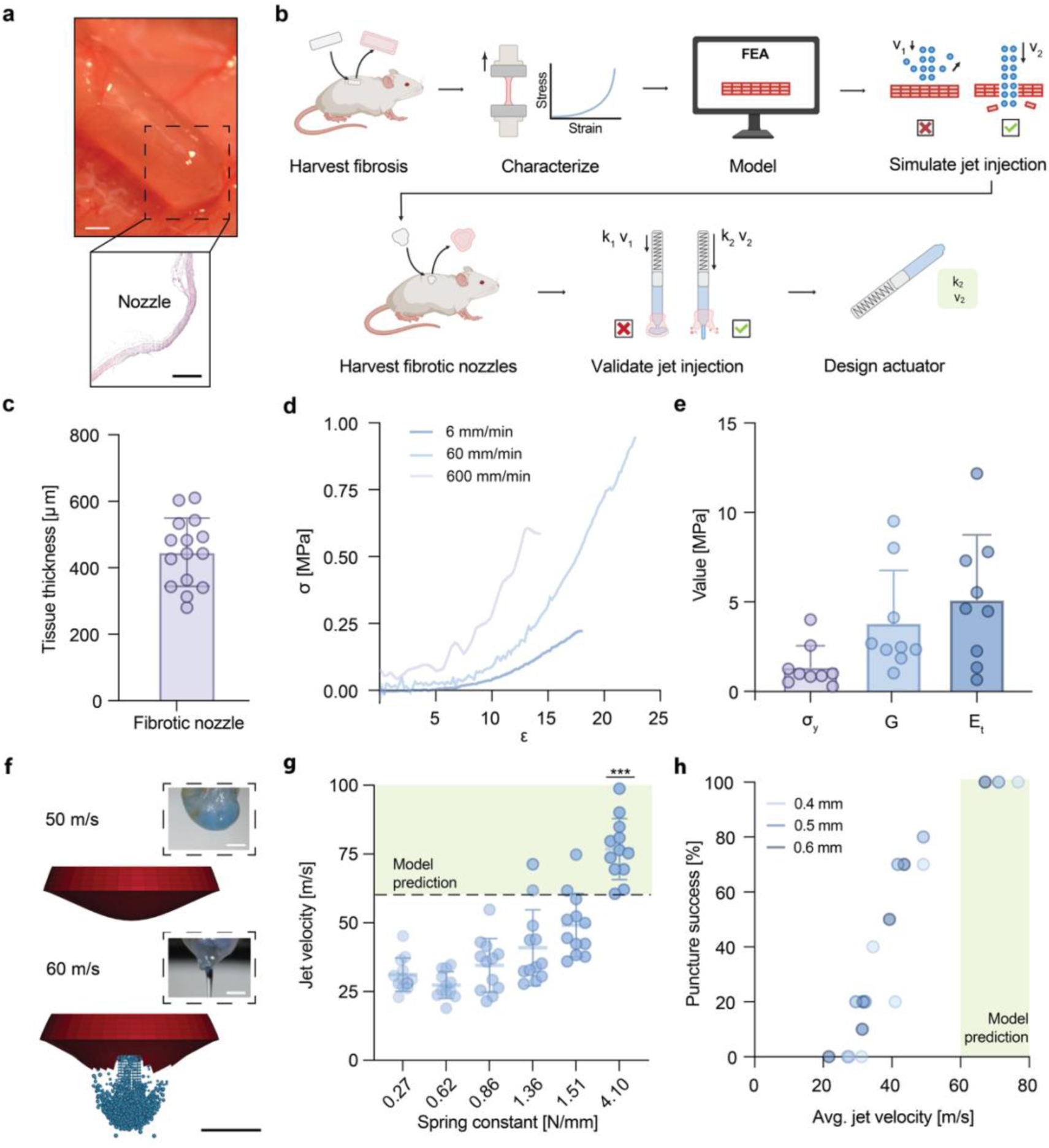
Mechanical characterization of jet injections puncturing fibrotic encapsulations. **a**, Optical (top) and hematoxylin and eosin (H&E) stained-histology (bottom) images of fibrotic encapsulation around implanted device (top scale bar, 3 mm; bottom scale bar, 300 μm) **b**, Experimental workflow characterizing jet injection through fibrosis. FEA = finite element analysis, v = velocity, k = spring constant. **c**, Thickness of fibrotic tissue around implanted nozzle. *n =* 15 nozzles. Data represent mean ± s.d and individual data points. **d,** Average stress (σ)- strain (ε) curves of rat fibrotic tissue at various testing speeds. *n* = 9 tissue samples averaged per group. **e,** Yield stress (σ_y_), shear modulus (G), and plastic hardening modulus (E_t_) from 60 mm/min testing speed. Data represent means ± s.d. *n* = 9 tissue samples per group, represented by dots. **f,** (Top) Representative image of jet failing to rupture fibrosis and model simulation of 50 m/s, ⌀ 0.4 mm jet impact on tissue. (Bottom) Representative image of successful jet rupture and model simulation of 60 m/s, ⌀ 0.4 mm jet impact on tissue (scale bars, 1 mm). The bottom inset image is repeated and annotated in figure 1g. **g**, Jet impact velocity through ⌀ 0.4 mm nozzle outlet across devices with springs of varying stiffness. *n*=12 devices per group. Data represent means ± s.d and individual data points. Line represents model prediction of fibrosis rupture at 60 m/s. Statistical significance of difference between jet velocities and the model prediction was computed via one sample, two tailed t-test. \*\*\**P* = 0.0003. This and additional jet velocity data can be found in Fig. S1. **h,** Success of varying jet velocities in puncturing ex vivo fibrotic tissue. *n*=10 tissue samples per group. Data points represent percent success. Illustrations in **b** created with BioRender.com.

We first subcutaneously implanted dummy devices in rats for a month and harvested fibrotic tissue to perform mechanical characterization. Fibrotic tissue around implanted nozzles measured 447 ± 103 µm (n=15), consistent with human fibrotic capsule thicknesses around similarly sized subcutaneous drug rod implants^30^ (Fig. 2c). Tensile testing on harvested fibrotic tissue revealed elastic and plastic stress-strain regions from which we extracted shear moduli, yield stresses, and plastic hardening moduli to inform our model (Fig. 2d-e). These results align with prior research characterizing the tensile strength of breast implant fibrosis^31^.

To simulate the jet-tissue interaction, we implemented a finite element method-smoothed particle hydrodynamics (FEM-SPH) approach that utilized our measured fibrosis material properties. Then, we evaluated the erosion of the tissue mesh with different jet diameters and initial velocities (Fig. 2f). Simulations using the average measured fibrosis material properties showed complete erosion of the tissue mesh at jet velocities ≥60 m/s for a jet diameter of 0.4 mm and jet velocities ≥50 m/s for jet diameters of 0.5 and 0.6 mm. A sensitivity analysis varying the fibrosis material properties from their minimum to maximum recorded values showed changes in predicted penetration velocities of up to +10 / −20 m/s (Table S3). These values are 30-40 m/s higher than velocities required for water-cutting other soft tissues, such as the liver^32^. Below these threshold velocities, the jet particles rebounded from the mesh surface.

*Ex vivo* jetting studies through excised fibrotic tissue validated model findings. To reproduce jet velocities from the simulation, we developed a manually triggerable jet injection device that accommodated springs and nozzles of varying stiffnesses and outlet diameters, respectively. Using different configurations, we achieved average jet velocities from 21 to 76 m/s (Fig. 2g, Fig. S1). The strongest spring punctured fibrotic tissue with 100% success (n=10 tissue samples) at 67-76 m/s across all nozzle outlet diameters, while the weakest spring at 21 m/s never successfully broke through the tissue (Figure 2h). This is consistent with our model prediction threshold of 50 to 60 +10/-20 m/s.

We performed additional *ex vivo* jetting studies on rat skin to evaluate the impact of these jet velocities on surrounding tissue. Hematoxylin and eosin (H&E) histology staining revealed that the strongest jet only penetrates the hypodermis (Fig. S2), indicating that the delivered drug would remain in the subcutaneous space following injection.

Combined, these studies suggest that a spring-based jet injection system enables liquid drug penetration through fibrotic encapsulation and into the vascularized subcutaneous tissue for rapid uptake.

### Engineering a triggerable, syringe-implantable, rapid-acting drug reservoir

Utilizing the springs and nozzles characterized above, we engineered a millimeter-scale, triggerable actuation mechanism capable of reliably penetrating through fibrotic encapsulation (Fig. 3a). In the device’s idle state, a dissolvable polyvinyl alcohol (PVA) film holds the spring in compression. When triggered, an electrolytic pump begins generating gas inside a closed chamber filled with electrolyte fluid, propelling the remaining electrolyte through a small channel towards the PVA film. As the film hydrates and weakens, the spring quickly releases and drives a plunger to eject liquid formulations from the reservoir. Compared to heat and pump-based actuation mechanisms, electrolytic pumps use orders of magnitude less energy and space (Table S4). The entire actuation mechanism measures 16.7 x 3 mm.

**Fig. 3:**
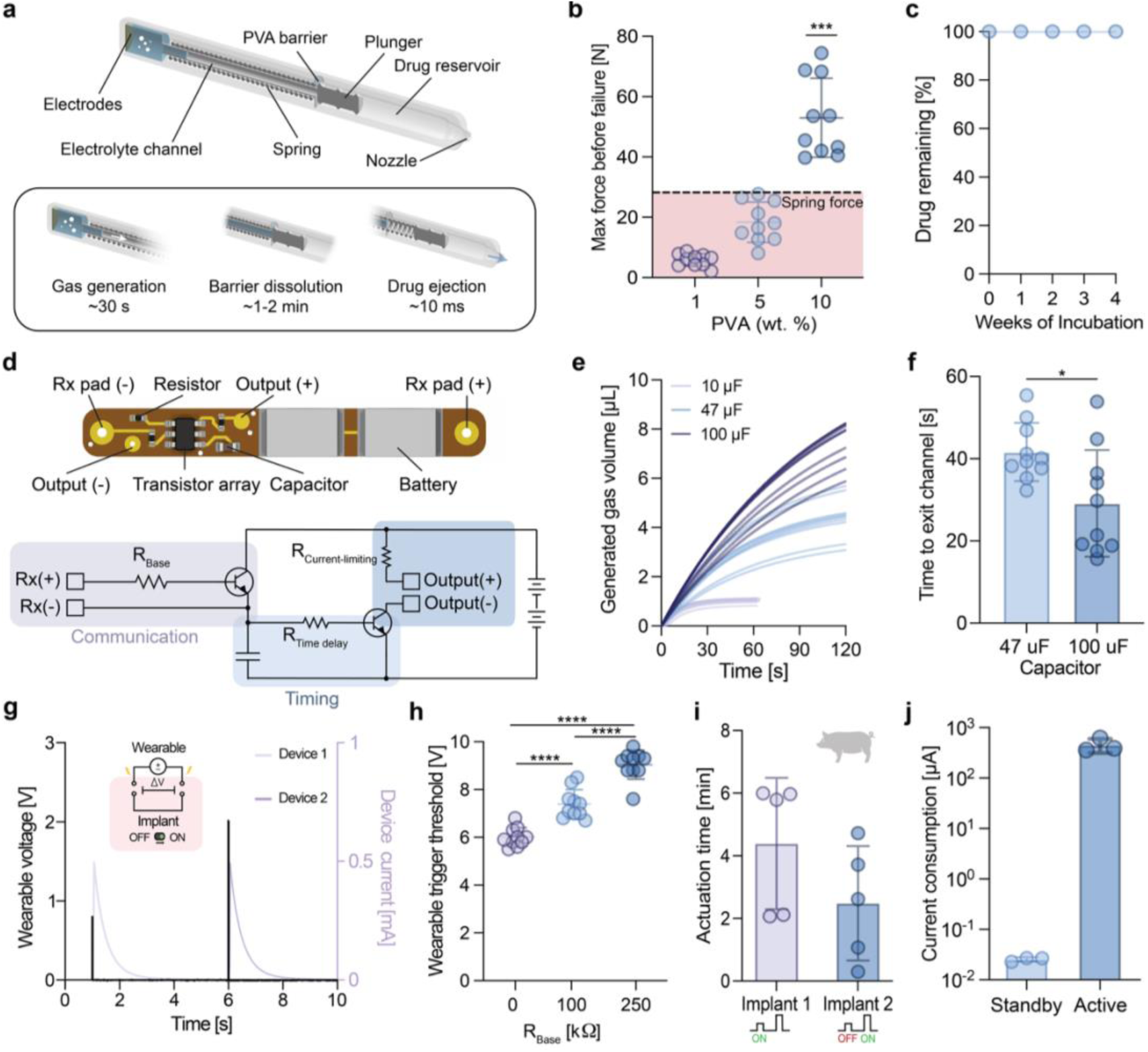
RES-Q implant design. **a,** Schematic of electrolytic and spring-based actuation mechanism for drug delivery. The device measures ⌀3 x 29.2 mm. **b**, Forces that polyvinyl alcohol (PVA) films prepared from solutions of varying weight (wt) % withstand before failure. *n*=10 films per group. Data represent means ± s.d and individual data points. Line represents the force that the strongest spring exerts in compression. Statistical significance of difference between maximum force before failure and the spring force was computed via one sample, two-tailed t-test. \*\*\**P* = 0.0002. **c,** Leakage test from devices with ⌀ 0.5 mm nozzle outlet. *n*=4 devices. No leakage was detected. Data represents mean drug remaining in implant ± s.d. **d**, Schematic of the flexible printed circuit (25 x 3 x 1.1 mm^3^) that powers the implant and enables triggering via in-body communication with the wearable monitor. Rx = receiver. R = resistor. **e**, Electrolytic gas generation propels the electrolyte towards the PVA barrier, and the pump rate changes with varying circuit capacitance. *n*=10 devices per group. Data represents individual trials. **f**, Time for the electrolyte to reach the PVA barrier through the channel with varying capacitors. *n*=10 devices per group. Data represent means ± s.d and individual data points. Statistical significance of difference between groups was computed via unpaired two-tailed t-test. \**P* = 0.0156. **g,** Representative benchtop demonstration of sequential implant triggering via distinct voltage pulse triggers from the wearable. Device 1 requires a lower voltage threshold to turn on than device 2. Data represents individual device trial. **h**, Implant circuits with different base resistances (R_Base_) require different triggering minimum voltage thresholds from the wearable, allowing sequential device triggering by moving from low to high voltage pulses. Data collected in ex vivo chicken breast tissue. The pulse from the wearable travels via tissue-mediated ionic conduction to the implants. n=10 locations in the tissue per group. Data represent means ± s.d and individual data points. Statistical significance of difference between groups was computed via ordinary one-way analysis of variance (ANOVA), F_(2,27)_ = 78.04 (****P <0.0001). **i.** Timing of sequential implant actuation in swine. Implant 2 was triggered after Implant 1 by increasing the voltage pulse applied to the tissue. *n = 10* devices in 2 swine. Data represent means ± s.d and individual data points. **j,** Max current consumption of RES-Q implant when actively waiting for an overdose signal (standby) and during electrolysis actuation (active). *n* = 3 implants. Data represent means ± s.d and individual data points. Illustrations in **g** and **i** created with BioRender.com.

The PVA film acts as a hydration-responsive barrier that maintains the spring in compression until weakened by fluid. PVA is ideal for its tunable mechanical properties and water solubility. We chose a high molecular weight (85,000-124,000), partially hydrolyzed (87-89%) PVA to provide compressive strength when dry and dissolution when wet at body temperature. Compression tests demonstrated that films prepared from 10 wt% solutions withstood a maximum force of 52 ± 13 N, 1.8x greater than the strongest spring’s exerted force in compression (Fig. 3b). Benchtop trials showed the film weakens to allow spring release 100 ± 30 seconds (n=10 devices) after fluid exposure, ensuring actuation within the narrow therapeutic window for overdose rescue.

A 150 µm thick polydimethylsiloxane (PDMS) valve at the nozzle tip prevents drug leakage prior to actuation through hydrophobic effects. Furthermore, a moisture impermeable, biocompatible ceramic coating around the actuation mechanism ensures prolonged actuator stability in a 37°C, high humidity environment (Fig. S3). Leakage studies performed with submerged drug reservoirs in a 37°C 50 RPM incubating shaker showed no detectable drug leakage over four weeks, supporting long-term device viability (Fig. 3c).

An accompanying syringe-implantable, flexible printed circuit board (fPCB) powers the electrolytic pump and enables communication with the wearable sensor hub (Fig. 3d). Upon overdose detection, the wearable hub initiates communication by emitting an electrical pulse into the tissue. Because body tissue is ionically conductive, tissue-interfacing receiving pads on the implanted fPCB detect this emitted pulse and trigger the implant to switch on the electrolysis reaction. These pads bias an NPN transistor, which turns on only if the detected voltage approaches its saturation region. As shown in our previous studies, this ensures that low voltage ambient noise cannot switch on the device^29^. Our previous studies also show that fPCBs implanted long-term continue to detect emitted pulses even when encapsulated by fibrosis up to 1 mm thick^29^. This tissue-mediated communication system requires a narrow, 3-mm wide electronic footprint, enabling a syringe-injectable design.

After the implant is triggered by a short pulse from the wearable, an on-board delay circuit maintains the implant in an “on” state until a capacitor in the circuit is fully discharged, allowing for a prolonged electrolytic pump action. This differs from our previous work where our in-body communication protocol^29^ required the implant to receive constant wearable stimulation to remain on, and it mitigates tissue damage from prolonged stimulation. By varying the capacitor, we can control the duration and rate of the electrolytic reaction. During benchtop trials, a 100 μF capacitor generated enough gas in 29.2 ± 13 seconds for fluid to exit the channel and reach the PVA film (Fig. 3e-f).

A key advantage of this platform is its capacity for on-demand repeat-dosing (Fig. 3g). While a single dose of naloxone is typically sufficient for rescue, 15-25% of individuals require a second dose^33^. Our communication protocol allows for selective communication with multiple implanted devices by interchanging a single resistor on the fPCBs. This resistor lies in series with a tissue-interfacing pad, increasing the saturation voltage required to switch the implant on. For example, the first implant can be triggered by delivering a 7V pulse from the wearable, while a second device can be triggered by delivering a 10V pulse from the wearable (Fig. 3h). With this distinct switching criteria, a second dose can be delivered if one dose is insufficient for rescue. We demonstrated this switching criteria and confirmed actuation of 10 of our fully assembled devices *in vivo* in swine (Fig. 3i, Fig. S4).

The syringe-implantable fPCB can support two rechargeable 3 x 4.4 x 1.1 mm^3^ 100 uAh TDK CeraCharge batteries or one ⌀ 3 x 12 mm^3^ 7 mAh CR311 battery. Due to the passive nature of our communication mechanism, the fPCB can remain in a standby state with a 25 nA average current draw (Fig. 3j). This enables theoretical ultra-long battery lifetimes prior to actuation of up to 5.5 months per charge using the TDK batteries or 30+ years using the CR311 battery. This can be compared to the one-to-two-week lifetimes per charge of other larger prototype closed-loop delivery systems^10, 11^. During electrolysis, the device consumes a maximum of 455 μA on average (Fig. 3j). With the ultra-low power and size requirements of our actuator and communication components, our fPCB has a footprint of either 3 x 1.1 x 25 mm^3^ or 3 x 3 x 27.5 mm^3^ with the CeraCharge or CR311 batteries, respectively.

### Closed loop overdose rescue

To demonstrate the closed-loop, sensor-mediated, drug delivery capabilities of the RES-Q network, we implanted devices in rats and subsequently induced and treated a fentanyl overdose. During an overdose, fentanyl leads to rapid respiratory depression and cardiovascular complications such as bradycardia and hypoxia^34, 35^. Unlike prior experimental devices which utilize on-board sensors to detect an overdose, we designed a two-part wearable (Fig. S5) and implant system (Fig. 4a) connected via tissue-mediated ionic communication. This system architecture dramatically reduces implant size to enable syringe-injectability and keeps power consumption low to extend battery lifetimes. Furthermore, the ultralong 40 cm communication range through tissue enabled by ionic communication allows the sensors and implants to be placed in disparate locations that maximize sensitivity, safety, and convenience^29^.

**Fig 4.**
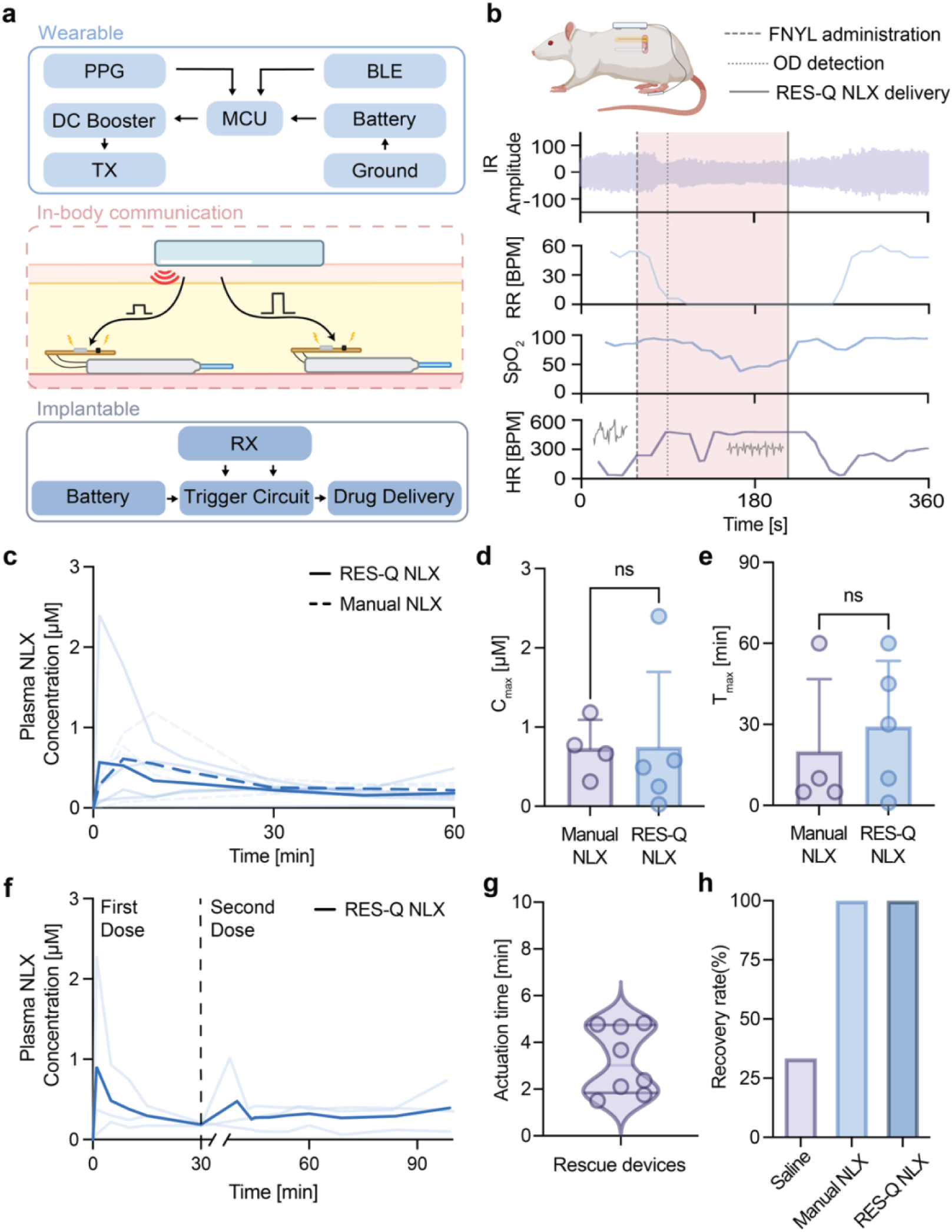
RES-Q platform-mediated closed-loop overdose rescue. **a,** Schematic of RES-Q platform for closed-loop control of implanted autoinjectors. PPG = photoplethysmography, DC = direct current, TX = transmitter, MCU = microcontroller unit, BLE = Bluetooth Low Energy, RX = receiver. **b,** Representative rat vital signs collected from RES-Q network during a fentanyl overdose and rescue. This and additional animal trial data can be found in Fig. S9. The inset traces in the heart rate (HR) graph represent infrared (IR) signals that denote a change in pulse amplitude following fentanyl (FYNL) administration. OD = overdose, NLX = naloxone, RR = respiratory rate, BPM = beats or breaths per minute. SpO_2_ = Peripheral capillary oxygen saturation. **c,** Pharmacokinetic profiles of naloxone following manual injection and RES-Q device delivery. *n* = 5 rats with RES-Q NLX. *n* = 4 rats with a manual NLX injection. Data represents mean (bold) and individual trials. **d**, Maximum plasma concentration (C_max_) for manual and RES-Q device delivery groups. *n* = 5 rats with RES-Q NLX. *n* = 4 rats with a manual NLX injection. Data represent means ± s.d. and individual data points. Statistical significance of difference between groups was computed via unpaired two-tailed t-test. P = 0.9782, not significant (ns). **e**, Time to maximum plasma concentration (t_max_) for manual and device delivery groups. *n* = 5 rats with RES-Q NLX. *n* = 4 rats with a manual NLX injection. Data represent means ± s.d. and individual data points. Statistical significance of difference between groups was computed via unpaired two-tailed t-test. P = 0.6058, not significant (ns). **f,** Pharmacokinetic profiles of sequential doses of naloxone from two RES-Q devices in the same rat. *n* = 3 rats. Data represents mean (bold) and individual trials. Time break represents 1–5-minute variations when preparing for second dose delivery. **g,** Time for RES-Q devices involved in rescue to deliver naloxone following overdose detection. *n* = 8 rats. Dots represent individual trials. **h,** Recovery rate for animals with a RES-Q device loaded with saline (*n* = 3 rats), a manual naloxone injection (*n* = 6 rats), and RES-Q devices loaded with naloxone (*n* = 8 rats). Illustrations in **a** and **b** created with BioRender.com.

The wearable hub (45 x 26 x 16 mm^3^) performs signal-quality-gated photoplethysmography (PPG) analysis to measure heart rate, pulse amplitude, and peripheral capillary oxygen saturation (SpO_2_) via infrared and visible light emission. On-device measurements and signal processing techniques were validated against simultaneously recorded electrocardiogram (ECG) and veterinary-grade sensor data, along with held-out reference recordings (Fig. S6). The onboard algorithm detects overdoses using continuous baseline changes in these metrics to autonomously trigger the implant (Fig S7, Materials and Methods). On average, the wearable draws 10.4 mA, corresponding to an estimated runtime of 28.9 hours of continuous monitoring per charge from a 300 mAh battery (Table S5). A single implant-triggering pulse draws <0.2 mWh from the batteries. We also demonstrate the capability to stream heart rate and blood oxygen levels from a commercial smartwatch to our RES-Q system as an alternative sensor input (Fig. S8).

The RES-Q network demonstrated timely therapeutic intervention following overdose detection. Animals exhibited signs of overdose within 45 ± 35 seconds (n=16) of fentanyl administration (15 µg/kg) as detected by our wearable. Although our device can hold a clinically relevant 0.4 mg human dose in our 16 µL reservoir at a concentration 25 mg/mL, we diluted our naloxone mixture to dose the rats with 1 mg/kg of naloxone. In a representative trial, (Fig. 4b), the wearable detected signs of an overdose 30 seconds after fentanyl administration. During this period, respiratory rate dropped from 64 to 6 breaths per minute (BPM), and arterial pulse amplitude dropped 30% from baseline. The reduction in pulse amplitude met the overdose detection criteria, prompting the wearable to initiate rescue. The device completed naloxone delivery 126 seconds after communication from the wearable and spontaneous respiration began 51 seconds later. Baseline respiratory rate was recovered 64 seconds after treatment. Additional detailed trial data from all tested animals can be found in the supplementary information (Fig. S9). After device triggering, blood plasma samples were collected to confirm naloxone uptake.

Pharmacokinetic evaluation demonstrated comparable naloxone uptake profiles between manual naloxone injections and RES-Q devices (Fig. 4c), including peak plasma concentration (C_max_) (Fig. 4d), time to peak plasma concentration (t_max_) (Fig. 4e), and area under the pharmacokinetic curve (AUC) (Fig. S10).

We also demonstrated sequential triggering of two separate implanted devices in the same animal, allowing for stepwise dosing. One implantable fPCB possessed a 0-ohm resistor in series with the receiving pad, while the second fPCB possessed a 215k-ohm resistor. The wearable delivered a 7V pulse upon overdose detection which triggered only the first implant. After thirty-five minutes, the wearable was manually prompted to deliver a 10 V pulse to trigger the second implant. Pharmacokinetic evaluation demonstrated sequential dose delivery (Fig. 4f). The AUC for the single dose and double dose RES-Q groups was 2.21 ± 2.37 µM·min and 4.91 ± 1.08 µM·min, respectively (Fig. S10).

Devices used for overdose rescue, in both the single dose and double dose groups, achieved naloxone delivery within the narrow therapeutic timeframe (3.21 ± 1.43 min, n=8) (Fig. 4g). All eight rats deployed with the RES-Q network loaded with naloxone and all five rats administered a manual naloxone injection recovered under anesthesia, while only one out of three rats deployed with a RES-Q device containing saline alone self-recovered (Fig. 4h).

## Discussion

In this study, we demonstrate triggerable, syringe-implantable, multi-dose drug delivery systems capable of overcoming the size and energy constraints associated with conventional sensing, communication, and actuation methods that have previously limited clinical applicability of closed-loop implantable devices. Fibrotic encapsulation of implantable drug delivery devices must be overcome to enable rapid uptake into the subcutaneous space. Here, we modelled the fracture mechanics of implant fibrosis to engineer an autoinjector mechanism with sufficient energy to puncture the tissue without being oversized or overpowered. Combined with our millimeter-scale electronics and in-body communication systems that replace bulky and energy intensive Bluetooth systems-on-chips, the autoinjector is small enough to fit within a 6-gauge needle and efficient enough to theoretically have a years-long battery lifetime. We demonstrate the RES-Q network’s ability to perform sensor-mediated, closed-loop delivery of both single and sequential doses of naloxone following a detected fentanyl overdose in rats.

To enable clinical translation, future overdose studies in large animal models and humans are required. Additionally, future studies quantifying the high-strain rate response of fibrosis and the material properties of human fibrosis could further improve the precision of our jet-injection model. Furthermore, years-long drug stability studies will be required to accurately assess naloxone stability in the RES-Q implant in vivo, although the drug maintains stability in environments that exceed body temperature^36^.

Clinical translation of the RES-Q network requires consideration of several key criteria. First, decoupling the sensor from the implant poses a risk that individuals will have poor adherence to a continuous wearable. While previous studies suggest that sensor-integrated watches or rings are accepted by patients for SpO_2_ and heart rate monitoring^37, 38^, future studies must assess multiple body locations on patients suffering from opioid addiction to identify the most comfortable area and device form factor that still provides adequate sensing and signal transmission to the implant. Additionally, the wearable’s day-long battery life necessitates occasional recharging, highlighting the need to investigate lower-power optical sensing and radio duty-cycling strategies to support sustained use. However, trading a longer implant battery lifetime for a shorter wearable device lifetime provides a simpler path for patient compliance through wall outlet recharging. Further work should also evaluate triggering specificity across broad physiological and ambulatory conditions; however, accidental naloxone dosing provides minimal side effects even during a false positive event^39^. Additionally, after triggering, removal of the device will require a small surgical procedure, but this process is well-established for small implants like Nexplanon^40^. Appropriate patient populations should be chosen that would benefit most from this device, such as those already on other addiction mitigation therapies.

While this study focuses on naloxone delivery, the RES-Q network is a platform technology that can load clinically relevant liquid doses of multiple emergency drugs, including epinephrine (Fig. S11). Glucagon and other biologics, however, would require a solid formulation to reach clinically relevant dose and stability levels. Additionally, our device can be deployed into the same in-body communication network as our previous implantable neuromodulation systems^29^ (Fig. S12), potentially allowing for sensor-mediated, cross-body, coordinated pharmacological and electrical interventions. Ultimately, the miniaturization and coordination between sensors and therapeutics achieved with this technology suggests that this method could enable patient-accepted devices for autonomous life-saving overdose rescue, and it justifies further evaluation for other diseases that require emergency intervention.

## Methods

### Implantable drug delivery device design and fabrication

The body of the drug delivery device was modeled with computer-aided design software (SOLIDWORKS 2025, Dassault Systèmes) and printed with biocompatible resin (BioMed Clear V2, Formlabs) using a three-dimensional (3D) printer (Formlabs 3B, Formlabs).

A ceramic coating (LR-100 LR Black, Cerakote) was added to the 3D-printed device components in some tests to reduce the material’s natural moisture permeability. Unless otherwise stated, the images of the device in the manuscript do not have the opaque ceramic coating in order to better show to components inside of the device. To prepare for coating, device parts were sprayed with isopropyl alcohol (IPA) and lightly sanded with 100-grit sandpaper. The ceramic coating was then applied to device parts using an airbrush at 25 psi until fully coated. Coated device parts were then cured in the oven at 180°C for an hour before resting at room-temperature for 24 hours.

Polyvinyl alcohol (PVA) film was prepared by dissolving PVA (Sigma-Aldrich, average M_w_ 85,000-124,000, 87%-89% hydrolyzed) in deionized water at 100°C while stirring for 6 hours. Ten milliliters of solution was cast into a 100 mm diameter Petri dish, allowed to cool at 40 °C, and cut into 2 mm diameter disks with a cutting machine (Silhouette Cameo 4) once solidified.

The 3D-printed nozzles had a polydimethylsiloxane (PDMS) inner coating prepared using a Sylgard 184 Silicone Elastomer Kit (Dow) with a 10:1 PDMS crosslinker ratio. The PDMS was applied with a needle to the inner surface of the nozzles and cured at room-temperature for 24 hours.

A schematic of the assembly process is shown in Fig. S13. First, PVA film was bonded to a rim inside the device body using medical grade epoxy (Loctite 4311). A full spring and an additional cut spring segment (CIM030A06 Compression Spring, Lee Springs) was then added to the device. The cut spring segment was added to achieve a maximum deflection that spans the full length of the drug reservoir. Next, the springs were compressed against the PVA film and secured with epoxy. A blunt 23-gauge needle, serving as the electrolyte channel, was inserted through the springs and epoxied. Then, a plunger, printed with a flexible biocompatible resin (BioMed Flex 80A, Formlabs), was coated in silicone oil (Super Lube, 5000 cSt) and inserted on the side of the PVA membrane opposite the springs. The device was then filled with 16 microliters of naloxone hydrochloride (Thermo Fisher Scientific) in 0.9% sodium chloride. After, electrodes were epoxied to the bottom of the electrolyte chamber, and the chamber was filled with 10 μL of 1 M NaCl. The chamber was then epoxied to the device. Wires were soldered from the output pads on the implantable flexible printed circuit board to the electrodes. The electrodes were designed with electronic computer-aided design (ECAD) software (KICAD 9.0) and outsourced for fabrication (JLCPCB). Lastly, the nozzle was epoxied to the drug reservoir.

Due to the manual, imperfect nature of device assembly throughout the experiments, nine out of the dozens of devices fabricated broke during testing. This occurrence is common amongst proof-of-concept device development studies. Results that arose from the broken devices were excluded from the data sets.

### Implantable fPCB design and fabrication

The implantable flexible printed circuit board (fPCB) was designed with ECAD software (KICAD 9.0) and outsourced for fabrication (JLCPCB). The circuit contains two gold-plated receiving pads (**⌀** 1.5 mm) that interface with the tissue and detect emitted voltage gradient signals. The presence of a voltage gradient switches an NPN transistor (MMDT3904) on, which activates a time delay circuit on the board consisting of an additional NPN transistor (MMDT3904), a capacitor (298W107X0004M2T, 100 µF 4V 0603), and a resistor (1 MΩ, 0201). Once the receiving pads detect a voltage gradient, current both flows to output pads on the board and charges the capacitor. When the voltage gradient is removed, the capacitor discharges through the second transistor. This transistor continues to conduct current through the output pads until the capacitor fully discharges. Modifying the capacitor and the resistor in the time delay circuit controls how long the output remains on. The board additionally has two areas for optional resistors. First, there is an area to add a resistor between the harvesting pad and the base of the first transistor. This increases the voltage necessary to switch the time delay circuit on, such that fPCBs with distinct switching criteria can be fabricated. In our closed-loop experiments, we fabricated boards with a 0-ohm resistor and boards with a 215k-ohm resistor in this location. Second, there is an area to add a resistor in series with the output pads to optionally limit the current through the output. In our experiments, we used a 0-ohm resistor here. Onboard footprints support 0201 surface mount resistors. The bare board measures 25 x 3 x 0.11 mm^3^, compatible with two 3 x 1.1 x 4.5 mm^3^ 1.5V solid state ceramic batteries (TDK CeraCharge). The final package measures 25 x 3 x 1.1 mm^3^. Alternatively, it is also possible to connect a CR311 battery, creating a final package size of 27.5 x 3 x 3 x mm^3^. The fPCB was encapsulated in medical grade epoxy (Loctite 4311) to isolate the electronics from the tissue, aside from the tissue-interfacing pads.

### Wearable hardware design and fabrication

The wearable consisted of a custom controller printed circuit board (PCB) and a separate photoplethysmography (PPG) sensor PCB (Fig. S5). The controller incorporated a XIAO nRF54L15 module (Seeed Studio, Shenzhen, China), which provided PPG acquisition, on-device processing, Bluetooth Low Energy (BLE) communication, and control of the implant-triggering circuitry. The system was powered by a rechargeable 3.7-V lithium-ion battery. The controller PCB supplied 3.3 V and 1.8 V to the PPG sensor through low-dropout regulators (TPS7A0233PDBVR and TPS7A0218PDBVR, Texas Instruments) and provided the I²C and interrupt interfaces.

PPG was acquired using a MAX30102EFD+T sensor (Analog Devices) mounted on the custom sensor PCB and connected to the nRF54L15 over I²C. During rat experiments, the sensor PCB was secured to the hind paw to record infrared and red PPG signals.

Implant-triggering voltage was generated using a TPS61088-based DC-DC boost converter. The nRF54L15 controlled the output voltage between 5 and 12 V using a pulse-width modulation (PWM) signal from GPIO D0, low-pass filtered and applied to the converter feedback input. A TPS22810DBVT load switch controlled by GPIO D1 electrically isolated the microneedle interface during standby and connected the programmed boost voltage only during an actuation pulse. The wearable electronics were packaged in a 3D-printed biocompatible resin enclosure (BioMed Clear V2, Formlabs) with medical adhesive (3M) for attachment to the skin.

### Microneedle fabrication

Microneedles were used to penetrate the skin and enable signal transmission into the tissue, as shown in our previous work^29^. In the same previous study, we also showed that conductive gel can be used to replace these microneedles. Stainless-steel shim (75 um, McMaster-Carr; 316SS) was cut with a femtosecond laser (60W, 75% power, 150 repetitions, or 4W, 60 kHz, 100% power, 45 repetitions, Optec) into microneedles. The microneedle prongs were raised from the cut shim with a razor blade and electropolished using a 6:3:1 glycerin: 85% phosphoric acid: DI water. Wire was lastly connected to each microneedle using conductive epoxy (CircuitWorks).

### Implantation procedure for harvesting fibrotic tissue

All animal procedures were approved by the Georgia Institute of Technology Institutional Animal Care and Use Committee and conducted in accordance with Georgia Institute of Technology animal facility guidelines (A100590). Adult male and female Sprague Dawley rats (200-350g; Charles River Laboratories, Wilmington, USA) were used for experiments. Animals were housed two per cage and maintained on a 12-hour light:12-hour dark cycle with a standard diet and continuous access to water.

Animals were anesthetized with isoflurane delivered in oxygen (5% induction, 2.25-3% maintenance) and maintained on a heated bed in ventral recumbency. Eye lubricant was then applied to both eyes with a cotton-tipped swab. The surgical site on the upper dorsum was shaved and prepared with three successive applications of alcohol and chlorhexidine solution in a circular pattern. A sterile fenestrated drape was placed over this surgical site.

A 2 cm incision was made through the skin on the dorsal side. Using blunt dissection, subcutaneous tissue pockets were created for device placement. Once devices were in place, the incision was closed using wound clips. Animals received a postoperative subcutaneous injection of ketoprofen (5 mg/kg) and sustained release buprenorphine (0.8 mg/kg) for pain management while recovering in a heated cage. Wound clips were removed 10-14 days after a procedure. After four weeks, animals were euthanized, and devices were explanted with surrounding fibrotic tissue intact.

### Thickness characterization of fibrotic nozzles

Nozzles (⌀3 mm x 2.7 mm) were modeled with computer-aided design software (SOLIDWORKS 2025, Dassault Systèmes, Waltham, MA) and 3D-printed with biocompatible resin (BioMed Clear V2, Formlabs). The nozzles were designed in connected sets of five (17.6 x 6 x 3 mm^3^) to more easily locate them during explantation. These nozzle sets were implanted subcutaneously in Sprague–Dawley rats for four weeks (Fig. S14). After four weeks, the devices were explanted with fibrotic tissue intact. The fibrotic tissue was later excised from each nozzle and measured with calipers. Three thickness measurements were taken along the tissue per sample and averaged per sample.

### Mechanical characterization of fibrotic tissue

Rectangular substrates (15 x 5 x 1 mm^3^) 3D-printed with biocompatible resin (BioMed Clear V2, FormLabs) were implanted subcutaneously in Sprague–Dawley rats for four weeks. After four weeks, the substrates were explanted with fibrotic tissue intact (Fig. S14) and immediately transferred to a specimen container of PBS where they were stored until use, within four hours of explanation. All mechanical characterization was performed using a tensile testing machine (Mark-10, ESM750) with a 10 N force gauge (Mark-10, Model M5-2). A custom data acquisition program (LabView) collected force and displacement data during tests. For tensile testing, the dimensions of the excised tissue were measured with calipers. The tissue was secured into grips with sandpaper to reduce slippage and stretched at 6, 60 and 600 mm/min until material failure. The resulting force versus displacement curves were converted to true stress versus strain curves. Shear modulus, yield stress, and plastic hardening modulus were extracted for input into the computational model. The shear modulus was approximated using Equation 1, where *E* is the Young’s modulus and ν is Poisson’s ratio.

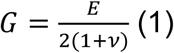

A Poisson’s ratio of 0.49, consistent with soft tissue, and Young’s modulus values determined by the slope in the linear regions of each curve were used for the approximation. Yield stresses were determined by the stress at the inflection point where the curves begin to exhibit plastic deformation. Lastly, the plastic hardening modulus was determined by the slope of the plastic region of the curves.

### Jet injection through fibrosis simulation

Simulations of the jet-tissue interaction were conducted in LSDYNA LS-Prepost 4.10 and LS-Run (ANSYS). The tissue was modeled as a three-dimensional 450 μm x ⌀3 mm cylinder, meshed with 2300 elements, and fixed in the y-direction around the edge. The mesh was assigned to an isotropic elastic material using the average shear modulus, yield stress, and plastic hardening modulus recorded during our mechanical characterization study. The density and bulk modulus of the tissue were approximated as those of water. The jet was then modeled as a cylindrical collection of particles. Eroding contact was defined between the particles and the mesh such that mesh elements were deleted if the impact met the maximum strain criterion, simulating tissue rupture. Erosion of the tissue mesh was evaluated across jet diameters of 0.4 mm, 0.5 mm, and 0.6 mm, and initial velocities of 10, 20, 30, 40, 50, 60, and 70 m/s. A sensitivity analysis was also carried out, where we assigned the mesh the entire range of material properties acquired during our studies and recorded the required jet velocity to penetrate the tissue.

### Jet parameters study

The effects of varying nozzle size and spring constant on jetting parameters were characterized with a simplified, spring-powered jetting apparatus. The apparatus consisted of a plunger and spring held in compression by a removable tab and a 16-microliter reservoir fitted with a nozzle. Pulling the tab of the apparatus released the spring, which drove the plunger to eject the liquid contents in the reservoir. A custom device to hold the jetting apparatus was 3D-printed with polylactic acid (PLA) filament using a fused deposition modeling (FDM) 3D-printer (Bambu Lab P1S, Bambu Lab) (Fig. S14). This holder was the clamped vertically above a 10-N force gauge (Mark-10, Model M5-2) with a compression plate attached. For each trial, an apparatus was loaded with dye and actuated while capturing the impinging jet with a high-speed camera (Edgertronic SC1). We simultaneously recorded the resulting maximum force on the compression plate using the force gauge. This was repeated with eighteen device iterations that varied in spring constant (CIM020A 07 S, CIM025A 07 S, CIM025A 04 S, CIM030A 07 S, CIM030A 06 S, LC 024A 09 S Compression Springs; Lee Springs) and nozzle hole diameter (0.4 mm, 0.5 mm, 0.6 mm). Springs were cut and added in series as needed to achieve a maximum deflection (free length − solid height) that spanned the full length of the drug reservoir (7 mm). The force of the jet impacting the gauge is given by Equation 2:

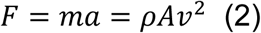

where *F* is the peak impact force, *m* is mass, *ρ* is the liquid density, *A* is the cross-sectional area of the jet, and *v* is the velocity. Rearranging Equation 2 gives the jet velocity in equation 3:

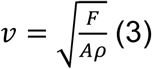

The kinetic energy (KE) of the fluid can be directly calculated from the mass (m) and velocity of the jet using Equation 4.

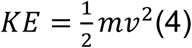

### *Ex vivo* jet injection study with fibrotic tissue

Nozzles (⌀3 mm x 2.7 mm) were modeled with computer-aided design software (SOLIDWORKS 2025, Dassault Systèmes, Waltham, MA) and 3D-printed with biocompatible resin (BioMed Clear V2, Formlabs). The nozzles were designed in connected sets (17.6 x 6 x 3 mm^3^) of five detachable nozzles. Three nozzle sets, one for each nozzle outlet diameter (0.4 mm, 0.5 mm, and 0.6 mm), were implanted subcutaneously in Sprague–Dawley rats for four weeks. After four weeks, the nozzle sets were explanted with fibrotic tissue intact (Fig. S14) and immediately transferred to a specimen container of PBS where they were stored until use, within four hours of explanation. Fibrotic tissue was then excised from each nozzle, and the fibrotic capsule was secured with glue to the jetting apparatus described in the jet parameter study (Fig. S14). For each trial, the apparatus was actuated, and success of puncture through the fibrotic tissue was recorded.

### Ex vivo jet injection study on rat skin

To evaluate the impact of a jet injection on surrounding tissue, skin was excised from freshly euthanized rats. The same spring-powered jetting apparatus and custom holder detailed in the Jet parameters study was used in this setup. The holder was clamped vertically 1 mm above the hypodermic side of a skin sample. An apparatus was loaded with tissue marking dye (CDI’s) and actuated. This was repeated with eighteen device iterations that varied in spring constant (0.27 N/mm, 0.62 N/mm, 0.86 N/mm, 1.36 N/mm, and 1.51 N/mm, 4.10 N/mm) and nozzle hole diameter (0.4 mm, 0.5 mm, 0.6 mm). Afterwards, the skin samples were fixed in 10% neutral buffered formalin (VWR) for histology. Tissue slides were prepared and stained with hematoxylin and eosin (H&E). The stained slides were analyzed using an optical microscope to visualize deposition of the dye.

### PVA strength study

Compressive failure of the PVA films was performed using a mechanical testing machine (Mark-10, ESM750) with a 10 N force gauge (Mark-10, Model M5-2). A custom data acquisition program (LabView) collected force data during tests. For each trial, PVA was adhered to the device was epoxy (Loctite 4311). The device was positioned between grips of the testing machine while the PVA was subjected to compression by a blunt 17-G needle connected to the force gauge. The maximum force prior to failure was recorded.

### Actuator stability study

Devices were assembled with a ceramic coating as detailed in the “Implantable drug delivery device design and fabrication” section and loaded with red dye in the drug reservoir.

The devices were then embedded in 1% agarose gel to mimic implantation in the subcutaneous space. Agarose gel was prepared by first dissolving agarose powder (Sigma-Aldrich) in deionized water. The solution was then poured into a mold and allowed to cool for two minutes before placing the devices. The gel was allowed to fully solidify for fifteen minutes before removal from the mold. The device-embedded gels were then placed in a glass Petri dish with a bath of phosphate-buffered saline (VWR) to keep the hydrogel hydrated. The dish was held in an incubating shaker at 37°C and 50 RPM, while the gels were monitored daily for any expulsion of red dye from the devices.

### Leakage study

Drug reservoirs with nozzle outlet diameters of 0.5 mm (n=4) were loaded with naloxone (25 mg/ml) and submerged in separate vials containing 4 mL of phosphate buffered saline (PBS). The vials were then placed in an incubated shaker set to 37 °C and 50 RPM for four weeks. On day 0, 7, 14, 21, and 28, 80 μL of PBS was sampled from the vials and replaced with 80 μL of fresh PBS. Sampled PBS was stored in microcentrifuge tubes and kept at −80 °C until analysis. On the day of analysis, samples were thawed and prepared for quantification with high-performance liquid chromatography (HPLC).^41^ Quantification was carried out with an Agilent 1100 Series modular HPLC system. Samples were injected in a mobile phase of methanol, acetonitrile and 50 mM phosphate buffer pumped at a flow rate of 1 ml/in through a Kromasil 100, C-18 (5 µ, 250 × 4.6 mm i.d.) column. Average retention time of naloxone was 11.3 minutes. For each device, triplicate injections were performed per sample, and the median value of the triplicates was used.

### Electrolysis gas generation and fluid travel study

The current supplied to the electrodes during electrolysis was calculated by measuring the voltage drop across a series 100-ohm resistor and applying Ohm’s Law. Data was collected with a tabletop multimeter (Keithley) and recorded using a custom LabView program, following the application of a 0.8 V 100 ms pulse to the harvesting pads of the circuit. This was repeated with a 10 µF, 47 µF, and 100 µF capacitor. During each experiment, the time for the electrolyte fluid to reach the end of the channel after the initiation of electrolysis was also recorded.

The moles of gas (*n*) generated at each electrode during electrolysis is given by equation 5:

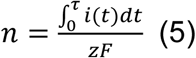

where *F* is the Faraday constant, equal to 96485 C/mol, *i* is current, *z* is the number of electrons required to obtain one gas molecule, and *t* is time. Assuming water electrolysis dominates and produces hydrogen and oxygen gas, the electrochemical reactions are given by:

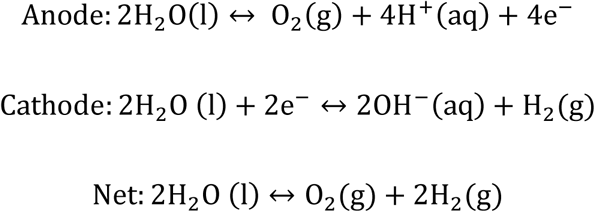

then

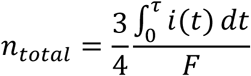

Substituting in the ideal gas law, the theoretical total volume (V) of gas generated is approximated in equation 6:

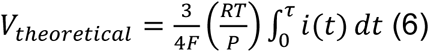

where R is the ideal gas constant, equal to 0.082507 L ⋅atm/(mol ⋅K), T is temperature, equal to 310.15 K, and P is pressure, equal to 1 atm.

### Benchtop demonstration of sequential implant triggering

To evaluate switching criteria, two fPCB circuits were replicated on a breadboard. A RES-Q device was connected to a circuit incorporating a 100-ohm current-limiting resistor, 0-ohm base resistor, and 0.47 µF capacitor, while another RES-Q device was wired to a circuit incorporating a 100-ohm current-limiting resistor, 22k-ohm base resistor, and 0.47 µF capacitor. A function generator (SDG2000X, Siglent) was then connected to the receiving pads of the circuit. An oscilloscope (SDS824X HD, Siglent) was additionally connected to the probes, and the function generator was programmed to deliver a 0.8 V pulse and a 2 V pulse. Voltage across the current-limiting resistors was collected with a tabletop multimeter (Keithley) and recorded using a custom LabView program, following the application of the pulses. The current supplied to the output of either device was calculated by applying Ohm’s Law.

### Transistor switching in *ex vivo* chicken breast

Chicken breast (Heritage Farms) was soaked in PBS (VWR) during the experiment. To evaluate switching criteria, the fPCB was replicated on a breadboard and the NPN transistor’s base resistor was switched out to be either 0-ohms, 100k ohms, or 250k ohms. Two 25-G hypodermic needles (Sol-M) served as the tissue-interfacing receiving pads and were placed at a fixed tissue separation of 22 mm in the chicken breast. Two additional 25-G hypodermic needles connected to a function generator (SDG2000X, Siglent) served as the voltage transmitting probes and were placed into the tissue 1 mm apart from the receiving pads. The applied voltage needed to turn on an LED was recorded for each resistor at the same location in the chicken breast. This was then repeated at ten different locations.

### Implant switching in in vivo swine study

All animal procedures were approved by the Georgia Institute of Technology Institutional Animal Care and Use Committee and conducted in accordance with Georgia Institute of Technology animal facility guidelines (A100022). Female Gottingen pigs (15-25 kg) were sedated, intubated, and anesthetized. The animals were placed in lateral recumbency, and an incision was made into the skin using a scalpel. Blunt dissection of the subcutaneous space was performed to create room for implants. Two implants were placed in the space: one with a 0-ohm resistor in series with the harvesting pad on its fPCB and another with a 215k-ohm resistor in series with the harvesting pad on its fPCB. The drug reservoirs were loaded with green or blue tissue marking dye (CDI’s) to enable histological assessment of the surrounding tissue after firing. Two 25-G hypodermic needles (Sol-M) were inserted into the skin on opposite ends of the incision and connected to a function generator (SDG2000X, Siglent). A 5 V pulse was delivered to activate the 0-ohm resistor implant, followed by a 10 V pulse to activate the 215k-ohm resistor implant one minute later. The time between the pulse delivery and implant firing was recorded. Implant firing was confirmed via auditory cues (click from the spring extension) and visual cues. Skin and fat with tissue marking dye was then excised and fixed in 10% neutral buffered formalin (VWR). Tissue slides were prepared, and a hematoxylin and eosin (H&E) stain was performed. The stained slides were analyzed using an optical microscope to visualize deposition of the dye. Swine were euthanized.

### Closed-loop opioid overdose rescue experiment

All animal procedures were approved by the Georgia Institute of Technology Institutional Animal Care and Use Committee and conducted in accordance with Georgia Institute of Technology animal facility guidelines (A100590). Rats were anesthetized with isoflurane delivered in oxygen (5% induction, 2.25-3% maintenance) and maintained on a heated bed. Tail vein and jugular catheters were placed for fentanyl administration and blood collection, respectively. All *in vivo* studies utilized a functional prototype of the final wearable board. The PPG sensor was secured to the rat’s hind paw with medical tape. For all animals, fentanyl citrate (15 ug/kg) was administered via the tail vein catheter after collecting baseline PPG data for 60 s.

Animals in the positive control treatment group received a manual subcutaneous injection of naloxone hydrochloride in 0.9% sodium chloride (16 µl of 1 mg/kg, Thermo Fisher Scientific) when the detection software indicated an overdose.

Animals in the negative control device group, single dose group, and double dose group were implanted with the RES-Q device(s). These animals were placed in lateral recumbency, and a region was shaved for microneedle placement. Microneedles were placed 22 mm apart and connected to the wearable prototype. A 6-gauge needle was inserted into the skin to create an opening for the implant, and surrounding subcutaneous tissue was bluntly dissected with scissors. The implant, loaded with naloxone hydrochloride in 0.9% sodium chloride (16 µl of 1 mg/kg, Thermo Fisher Scientific) for the treatment dose groups or solely 0.9% sodium chloride for the negative control device groups, was then placed. After fentanyl administration, the time between the overdose-detection/wearable-signal-transmission and the firing of the implant was recorded. Implant firing was confirmed via auditory cues (click from the spring extension) and visual cues. For the double dose group, the second dose was initiated by a pulse from the wearable 35 minutes after the first dose. For some trials, respiratory rate was also collected using a MAX30102 sensor placed on the rat’s side to detect expansive and contractive movement associated with breathing. Blood was sampled through the jugular catheter before naloxone or saline delivery (t=0), and t = 1 min, 5 min, 10 min, 15 min, 30 min, 45 min, and 60 min after delivery. The samples were collected in EDTA plasma tubes (BD K_2_EDTA Microtainer) and stored on ice prior to centrifugation.

### Naloxone pharmacokinetic evaluation

Blood samples were centrifuged at 1,500g for 10 minutes to isolate plasma. The plasma was stored in microcentrifuge tubes and kept at −80 °C until analysis. On the day of analysis, samples were thawed and prepared for quantification with liquid chromatography-mass spectrometry (LC-MS). For sample preparation, 20 µl of plasma was mixed with 400 µL of LC-MS grade-methanol. Samples were then vortexed, centrifuged, and transferred to LC vials for analysis. Samples were injected (1 µL) in two mobile phases, A: water/ 10mM ammonium acetate/ 0.1% acetic acid and B: 80% isopropanol/ 20% acetonitrile/ 0.1% acetic acid. The column (Waters Cortecs T3, 100 mm x 2.1 mm, 1.6-micron particle) was maintained at 45 °C. A mass spectrometer (IQ-X) was operated with a scan range of 100-1000m/z and full scan resolution of 60000.

One of the pharmacokinetic curves from the subcutaneous delivery of naloxone had a contaminated non-zero concentration of naloxone at t=0 minutes. This curve was excluded from the pharmacokinetics analysis and likely arose due to sample contamination at the analysis phase.

### Sensor data handling

All PPG signal preprocessing, peak detection, heart rate estimation, and boost converter control were executed on the nRF54L15, which closed the physiological loop on-device. These on-device algorithms are described in the following section. The host graphical user interface (GUI) application displayed the streamed data and supplied operator inputs (the boost-converter output-voltage setting and the drug-injection event), while the actuation decision itself was computed on-device. The on-device data path was sensor → I²C → nRF54L15 → BLE → host GUI.

The MAX30102 was configured once at start-up to drive the red (660 nm) and infrared (880 nm) light-emitting diodes (LEDs) in two-channel mode, sampling the red and infrared channels at 200Hz with 18-bit resolution (411 µs pulse width), an analog-to-digital converter range of 4,096 nA, and a drive current of 9.4 mA per LED. Samples were buffered in the 32-sample first-in-first-out (FIFO) register of the sensor with rollover enabled and FIFO sample averaging disabled, so that the output data rate equaled the 200 Hz sampling rate, and were read over I²C at a bus clock of 100 kHz, reduced from the 400 kHz default to increase signal margin. Because blocking I²C transactions cannot execute in interrupt context, acquisition used a deferred-work scheme in which a 5 ms (200 Hz) timer interrupt scheduled a work-queue task to perform the I²C and BLE operations. In each cycle, one red and one infrared sample were retrieved via a six-byte burst read, reconstructed to 18-bit values, and a millisecond timestamp from the device uptime counter was recorded and expressed relative to the measurement start time. Acquiring one sample pair per cycle matched the 200 Hz polling rate to the sensor’s 200 samples s⁻¹ output.

Data were streamed to the host via the Notify characteristic of the Nordic UART Service, gated on an active measurement state and a host subscription to the characteristic. The same link conveyed two host-to-device inputs: the operator-set boost-converter output-voltage parameter and the drug-injection marker. A connection interval of 7.5ms (peripheral latency 0, supervision timeout 4 s) was requested, and the peripheral was configured for an Attribute Protocol maximum transmission unit (ATT MTU) of 251 bytes. Each sample was serialized as one newline-terminated, comma-separated ASCII record containing the relative timestamp (ms), the raw infrared and raw red values (18-bit unsigned), the preprocessed infrared and red values (signed), the contact state, the heart rate (beats min⁻¹), the beat count, and the boost-converter on/off state. The heart rate and beat count were computed once per analysis window (10 s) and carried forward in every record until the next window completed. They were left empty during the first window, before any estimate was available. The host logged the received records to a comma-separated value (CSV) file, one column per field, for offline analysis.

### On-device PPG processing and heart-rate estimation

Red and infrared PPG signals were acquired at 200 samples s⁻¹ and processed locally on the nRF54L15 in non-overlapping 10-s windows. Each window therefore contained 2,000 samples per optical channel. For physiological analysis, the mean was removed from the infrared signal, and a zero-phase third-order Butterworth band-pass filter (2-20 Hz) was applied. Pulse amplitude was defined as the peak-to-peak range of the filtered infrared waveform. Heart rate was estimated from the autocorrelation of the filtered infrared PPG signal. The dominant autocorrelation peak within the predefined physiological heart-rate range was identified, and its lag was converted to heart rate according to equation 7:

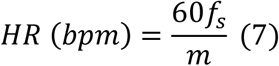

Where *f_s_* = 200 Hz is the sampling rate and *m* is the median inter-peak interval in samples. To prevent unreliable heart-rate estimates from entering the closed-loop decision, each window underwent an on-device heart-rate quality check. Rhythm strength was quantified from the magnitude of the selected autocorrelation peak, and a beat-doubling ratio was used to identify cases in which the estimated period corresponded to twice the true beat interval. Heart-rate estimates were accepted when the rhythm strength was ≥0.30 and the beat-doubling ratio was <0.70. If either criterion failed, heart rate was excluded from the overdose decision for that window, while the infrared pulse-amplitude measurement remained available for evaluation. The embedded implementation reproduced reference beat locations and quality decisions (Fig. S15).

### Closed-loop overdose detection and actuation

The closed-loop control algorithm was executed entirely on the nRF54L15. Before drug administration, heart rate and infrared pulse amplitude were monitored to establish subject-specific physiological baselines. At the operator-marked drug-injection event, the median heart rate and median infrared pulse amplitude measured during the preceding baseline period were stored as reference values, and autonomous overdose monitoring was enabled.

For each subsequent 10-s window, the current heart rate and infrared pulse amplitude were compared with their respective baseline values. An overdose condition was detected when either the quality-approved heart rate fell below 40% of its baseline value or the infrared pulse amplitude fell below 70% of its baseline value. Because heart rate was used only when its quality criteria were satisfied, a rejected heart-rate estimate did not independently trigger or suppress the amplitude-based decision.

When an overdose condition was detected, the nRF54L15 controlled the boost-converter and load-switch circuitry to deliver a stimulation pulse to the implant at the operator-configured output voltage and pulse duration. Following each actuation, an automatic shut-off terminated the stimulation pulse, and a refractory period prevented repeated actuation. The boost converter could also be activated manually from the host interface when required.

Because the closed-loop decision was executed locally, implant triggering was maintained during simulated BLE interruption (Fig. S16).

### Host communication and monitoring

BLE was used for configuration, visualization, and experimental logging but was not required for the closed-loop decision itself. The host interface provided operator inputs, including the drug-injection marker and stimulation settings, and displayed physiological measurements and device status. Signal preprocessing, heart-rate estimation, quality assessment, overdose detection, and actuation control were performed locally on the wearable nRF54L15.

### Smartwatch integration

A commercial smartwatch (Samsung Galaxy Watch 8 (Samsung Electronics, Suwon, Republic of Korea) running Wear OS) was integrated with the RES-Q platform to demonstrate compatibility with consumer wearable sensors. Heart rate and SpO₂ measurements from the watch were accessed using a custom Wear OS application developed in Kotlin and transmitted over a bidirectional BLE link to an nRF54L15-based receiver. The received physiological data were then passed to the RES-Q wearable platform for downstream processing and implant triggering.

### Validation of on-device PPG processing

On-device heart-rate estimates were evaluated using held-out recordings not used for algorithm development and compared with an offline reference. A subset was additionally compared with simultaneously recorded electrocardiogram (ECG)-derived heart rate or heart rate and SpO_2_ from a veterinary grade monitor (SurgiVet) (Fig. S6). Numerical fidelity of the embedded implementation was assessed by comparing the 32-bit nRF54L15 implementation with the corresponding 64-bit reference implementation.

### Wearable power consumption and battery life

Wearable power consumption was estimated in two firmware operating modes (Fig. S17). In the on-device processing mode used for the closed-loop experiments, PPG preprocessing, heart-rate estimation, quality assessment, overdose detection, and actuation control were executed locally on the nRF54L15, while the host remained connected for monitoring and logging. In on-device processing mode, the wearable generated one compact result summary per 10-s window (approximately 15 B s⁻¹ and 0.1 messages s⁻¹). In comparison, raw-streaming mode generated 200 ASCII records s⁻¹ (approximately 4.7 kB s⁻¹), of which 101 records s⁻¹ (approximately 2.3 kB s⁻¹) were received by the host during the measured session. Unreceived records were dropped and not retransmitted. Thus, the reported reductions in data volume (>300-fold) and message count (2,000-fold) refer to application-level traffic generated by the firmware rather than successfully delivered BLE throughput. These rates exclude BLE protocol overhead. Average current was estimated from the battery discharge trajectory rather than measured directly with a current meter. A 300-mAh lithium-polymer cell was charged until the charge indicator extinguished and the terminal voltage was stable at 4.15V. The terminal voltage was recorded at the battery pads with a handheld multimeter (10-mV resolution) at intervals of 30-60 min. One session was recorded per mode from the same cell, and the same charge-termination point at 25 to 26 °C. The charge consumed over each session was obtained from the difference in state of charge between the first and last reading, read from a generic lithium-polymer open-circuit-voltage curve and scaled by the nominal cell capacity, and the session-average current as that charge divided by the session duration. The energy per analysis window is that current multiplied by nominal cell voltage (3.7V) and the 10-s analysis window, and battery life is the nominal cell capacity divided by the current. Throughout each session, a host held the Bluetooth connection open and logged the telemetry, from which the fraction of the session with the link connected was obtained. The per-window computation time was obtained from the on-device session, where the firmware reports it for every window, and reflects the flashed build, which evaluates all three candidate peak detectors.

The optical sensing current in Table S5 is the sum of the average LED current and the supply current of the MAX30102. Neither was measured separately. The average LED current is NLED × ILED × PW × fs (two LEDs, 9.4 mA, 411 µs, 200 Hz: 1.55 mA). The supply current is the datasheet value (0.6 mA typical, 1.2 mA maximum). The current of the nRF54L15 was taken from the datasheet (CPU active, 2.6 mA; System ON idle, 2.9 µA; radio receive, 3.4 mA; radio transmit, 4.8 mA at 0 dBm and 9.8 mA at +8 dBm; 3.0 V, DC/DC regulator enabled). The duty cycle is 12 percent in raw streaming and 7.9 percent in on-device processing, combining an assumed 0.25 ms of acquisition work per 5 ms sampling cycle with either record formatting or window processing. The time the device spends computing one heart-rate value was measured on the device with a 1-us timer over 3,552 windows. It is 247 ms per 10-s window for the firmware as flashed, which evaluates all three candidate detectors, and 119 ms when only the selected detector is run.

### Software

A custom GUI was developed in Python (PySide6; The Qt Company, Espoo, Finland) for communication with the nRF54L15. The GUI provided two operating modes: real-time monitoring and offline analysis of previously recorded data.

In real-time mode, an output filename was specified, and a BLE connection to the nRF54L15 was established. The PPG Real-time Monitor window displayed, in real time, the raw IR and red signals, the window-wise preprocessed IR and red signals, the heart rate, SpO_2_, and the boost-converter output. The interface included controls for adding drug-event markers, the booster automatic/manual operation mode selection, and manually activating the boost converter. At the end of each session, the acquired data were automatically saved as CSV files.

In offline analysis mode, previously recorded signals were loaded and processed. The user could select and configure the procedures and parameters for preprocessing, peak detection, and SpO_2_ and heart rate calculation, and the results of each processing step were saved as a CSV file for subsequent review. A manual peak-annotation function allowed the user to mark peaks directly, updating the reference maps used by the adaptive peak-detection algorithm.

### Post-processing of closed-loop experiment data

Post-processed data from the closed-loop experiments can be found in Fig. S9. IR amplitude was found using the custom GUI’s offline analysis mode. On-device heart rate measurements collected during experiments were smoothed with a moving average using a window size of 3000. SpO_2_ measurements were calculating using the following equations 8 and 9^42^:

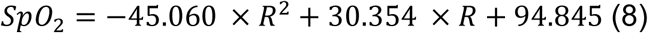

where

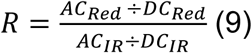

Variables in Equation 9 refer to the AC and DC component of the PPG waveform for red and infrared light. SpO_2_ measurements were then smoothed with a moving average using a window size of 6000.

### Neurostimulator implant and RES-Q implant triggering in *ex vivo* chicken breast

Chicken breast (Heritage Farms) was soaked in PBS (VWR) during the experiment. To evaluate switching criteria, two fPCB circuits were replicated on a breadboard. A RES-Q implant was connected to a circuit incorporating a 100-ohm current-limiting resistor, 0-ohm base resistor, and 0.47 µF capacitor, while a nerve cuff (NC-2-2-125SS-1.5-2-00-300-00, MicroProbes) was wired to a circuit incorporating a 100-ohm current-limiting resistor, 4.4k-ohm base resistor, and 0.47 µF capacitor. Two 25-G hypodermic needles (Sol-M) served as the tissue-interfacing receiving pads and were placed at a fixed tissue separation of 22 mm in the chicken breast. Two additional 25-G hypodermic needles connected to a function generator (SDG2000X, Siglent) served as the voltage transmitting probes and were placed into the tissue 1 mm apart from the receiving pads. An oscilloscope (SDS824X HD, Siglent) was connected to the transmitting probes, and the function generator was programmed to deliver a 7V pulse and a 10 V pulse. Voltage across the current-limiting resistors was collected with a tabletop multimeter (Keithley) and recorded using a custom LabView program, following the application of the pulses. The current supplied to the output of either device was calculated by applying Ohm’s Law.

### Statistical analysis

Experimental numbers and tests for all studies are indicated in figure legends. Statistical analysis was performed in GraphPad Prism Version 9 and 11. Statistical significance was based on measured P <0.05.

## Acknowledgements

We are grateful to all members of the Abramson Lab for helpful discussions. We thank R. Noel and L. O’Farrell for help with animal experiment design. We thank A. Asberry and D. Alexander with technical support in Georgia Institute of Technology’s Histology Core Facility. We thank the Parker H. Petit Institute for Bioengineering and Bioscience, Wallace H. Coulter Department of Biomedical Engineering, School of Chemistry and Biochemistry, and the Office of the Executive Vice President for Research for their generous support. This work was also supported by the Georgia Institute of Technology’s Systems Mass Spectrometry Core Facility.

## Author Contributions

J.J. and A.A. conceptualized and designed the experiments supporting the development of the RES-Q network. W.S. and J.M. developed the system software and communication interfaces for wearable device integration and external data transmission. J.J. developed the RES-Q device’s design. J.J., R.G., L.T., M.Aam., M.Ang., A.C., and S.H. fabricated the RES-Q device’s components. J.J., R.G., L.T., J.P., M.Aam., and M.Ang. performed benchtop and ex-vivo experiments for device characterization. J.J., S.H. Z.L, J.C., J.Z.D., and A.A. assisted with experiments in swine. A.A. and Y.J.L. provided funding and jointly supervised the project. All authors contributed to writing and editing the manuscript.

