## Supplementary Materials for "A network of syringe-injectable bioelectronics for closed-loop therapeutic rescue"

**The PDF file includes:**

**Tables S1 to S5**

**Figs. S1 to S17**

**Other Supplementary Material for this manuscript includes the following:**

**Source Data Fig. 2**

**Source Data Fig. 3**

**Source Data Fig. 4**

**Source Data Supplementary Figures**

### Table of Contents

**Supplementary Tables**

*Table S1. Comparison between recently reported techniques for controlling drug delivery.*

*Table S2. Comparison of implantable devices that demonstrate closed loop overdose rescue.*

*Table S3. Sensitivity analysis for the impact of mechanical tissue parameters on jet injection success in our finite element method-smoothed particle hydrodynamics model.*

*Table S4. Comparison of space and energy constraints for actuation mechanisms that enable triggerable drug delivery.*

*Table S5. Power budget and battery life of the RES-Q wearable by operating mode.*

**Supplementary Figures**

*Fig. S1. Jet velocities across different device iterations.*

*Fig. S2. Histological safety examination of jet injection on ex vivo rat skin.*

*Fig. S3. Actuator stability.*

*Fig. S4. Jet injection and histological safety examination in swine.*

*Fig. S5. Printed circuit board layout of the wearable sensing and voltage booster system*

*Fig. S6. Agreement between on-device and reference PPG processing.*

*Fig. S7. Overdose detection algorithm and triggering workflow.*

*Fig. S8. RES-Q platform integration with a smartwatch.*

*Fig. S9. Rescue data.*

*Fig. S10. Area under the curve (AUC) for the groups receiving naloxone.*

*Fig. S11. RES-Q device loaded with liquid epinephrine.*

*Fig. S12. Sequential in-body communication triggering of the RES-Q implant and the Smart Wireless Autonomous Networking System (SWANS) neurostimulator implant.*

*Fig. S13. Device assembly.*

*Fig. S14. Devices for fibrosis harvesting.*

*Fig S15. Agreement between on-device and reference photoplethysmography (PPG) processing.*

*Fig S16. Robustness of autonomous triggering to Bluetooth link interruption.*

*Fig. S17. Estimated wearable power consumption and nominal runtime in two operating modes.*

**Data Files**

Source Data Fig. 2. *Data included in Figure 2.*

Source Data Fig. 3. *Data included in Figure 3.*

Source Data Fig. 4. *Data included in Figure 4.*

Source Data Supplementary Figures. *Data included in Supplementary Figures.*

**<u>Supplementary Tables (Tables S1-S5)</u>**

| Reference | Type | Burst release | Premature leakage | Sensor-driven control |
|---|---|---|---|---|
| Ours | Autoinjector | Yes | No | Yes |
| *28* | Microsphere | Yes | No | No |
| *27* | Responsive hydrogel | No | No | No |
| *24* | Cellular implant | No | No | No |
| *25* | Cellular implant | No | - | No |

**Table S1. Comparison between recently reported techniques for controlling drug delivery.**

| Reference | Volume | Theoretical battery life | Titratable dosing? | Overcomes fibrosis? |
|---|---|---|---|---|
| Ours | 0.289 $cm^3$ | 30 years | Yes | Yes |
| [10] | 7.488 $cm^3$ | 1-11 weeks/charge | Yes | Yes |
| [11] | 18.564 $cm^3$ | 2 weeks/ charge | No | No |

**Table S2. Comparison of implantable devices that demonstrate closed loop overdose rescue.**

| Material property | Change in velocity required to penetrate relative to baseline [m/s] | |
|---|---|---|
| **Jet Diameter: 0.4 mm** | | |
| Yield Strain (Min/Max) | -10 | +10 |
| Yield Stress (Min/Max) | -10 | +10 |
| Shear Modulus (Min/Max) | -20 | +10 |
| Plastic Hardening Modulus (Min/Max) | -10 | 0 |
| **Jet Diameter: 0.5 mm** | | |
| Yield Strain (Min/Max) | -10 | +10 |
| Yield Stress (Min/Max) | 0 | +10 |
| Shear Modulus (Min/Max) | -10 | +10 |
| Plastic Hardening Modulus (Min/Max) | -10 | +10 |
| **Jet Diameter: 0.6 mm** | | |
| Yield Strain (Min/Max) | -10 | +10 |
| Yield Stress (Min/Max) | 0 | +10 |
| Shear Modulus (Min/Max) | -10 | +10 |
| Plastic Hardening Modulus (Min/Max) | -10 | 0 |

**Table S3. Sensitivity analysis for the impact of mechanical tissue parameters on jet injection success in our finite element method-smoothed particle hydrodynamics model**. The middle column represents the change in velocity required to penetrate the tissue between the minimum recorded material property value and the baseline average measurement. The right column represents the change in velocity required to penetrate the tissue between the maximum recorded material property value and the baseline average measurement.

| Reference | Actuation mechanism | Actuation size [$mm^3$] | Max power consumption [mW] | Breaks through fibrosis? |
|---|---|---|---|---|
| Ours | Electrolytic pump | 118.05 | 1.4 | Yes |
| [43] | Electrolytic pump | 57 | 4 | No |
| [44] | Electrolytic pump | 4 | 1 | No |
| [45] | Electrolytic pump | 56 | 1 | No |
| [46] | Mechanical pump | 469 | 252 | No |
| [11] | Mechanical pump | 537 | 132 | No |
| [47] | Mechanical pump | 424 | 7100 | No |
| [48] | Heater | 0.47 | 81 | No |
| [49] | Heater | 13 | 6000 | No |
| [50] | Heater | 495 | 225 | No |

**Table S4. Comparison of space and energy constraints for actuation mechanisms that enable triggerable drug delivery.**

| Mode | Raw streaming | On-device processing |
|---|---|---|
| Processing location | Host | Wearable |
| Messages produced ($s^{-1}$) / packets carried ($s^{-1}$) | 200 / 101 | 0.1 / 0.1 |
| Data sent ($Bs^{-1}$) | 4,700 | 15 |
| Connection interval (ms), requested by the firmware / granted by the host | 7.5 / 30 | 7.5 / 30 |
| **Power** | **Raw streaming** | **On-device processing** |
| Optical sensing current, LEDs + sensor (mA) | 2.2–2.7 | 2.2–2.7 |
| Microcontroller current, non-radio (mA) | 0.41 | 0.31 |
| Average battery current (mA) | 11.72 | 10.40 |
| Average power at 3.7 V (mW) | 43.4 | 38.5 |
| Energy per 10-s analysis window (mJ) | 433.6 | 3844.8 |
| Daily energy (mAh) | 280 | 251 |
| Daily energy (mWh) | 1040 | 923 |
| Battery life, 300-mAh cell (h) | 25.6 | 28.9 |
| Energy per implant-triggering pulse (mWh) | 0.08/pulse | 0.08/pulse |

**Table S5. Power budget and battery life of the RES-Q wearable by operating mode.** Both sessions used the same 300-mAh cell, charged to the same termination point and then discharged while its terminal voltage was read by hand with a multimeter. State of charge was converted from open-circuit voltage with a generic lithium-polymer table. Average battery current, average power, energy per analysis window, daily energy, and battery life all follow from those two voltage trajectories.

### Supplementary Figures

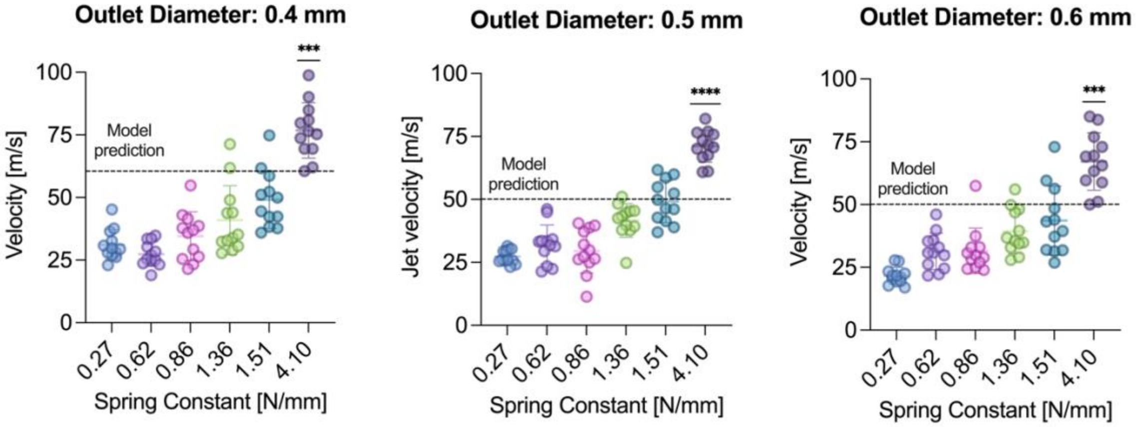


**Fig. S1. Jet velocities across different device iterations.** Jet impact velocities across three nozzle outlet diameters and six spring constants**.** The model prediction of 60 m/s is denoted for the 0.4 mm outlet diameter. The model prediction of 50 m/s is denoted for the 0.5- and 0.6-mm outlet diameters. The 0.4 mm outlet diameter data can also be found in Figure 2g and is presented again here for context. *n*=12 devices per group. Data represent means ± s.d and individual data points. Statistical significance of difference between jet velocities and the model prediction was computed via one sample, two tailed t-test. ***$P$ = 0.0003, ****P < 0.0001

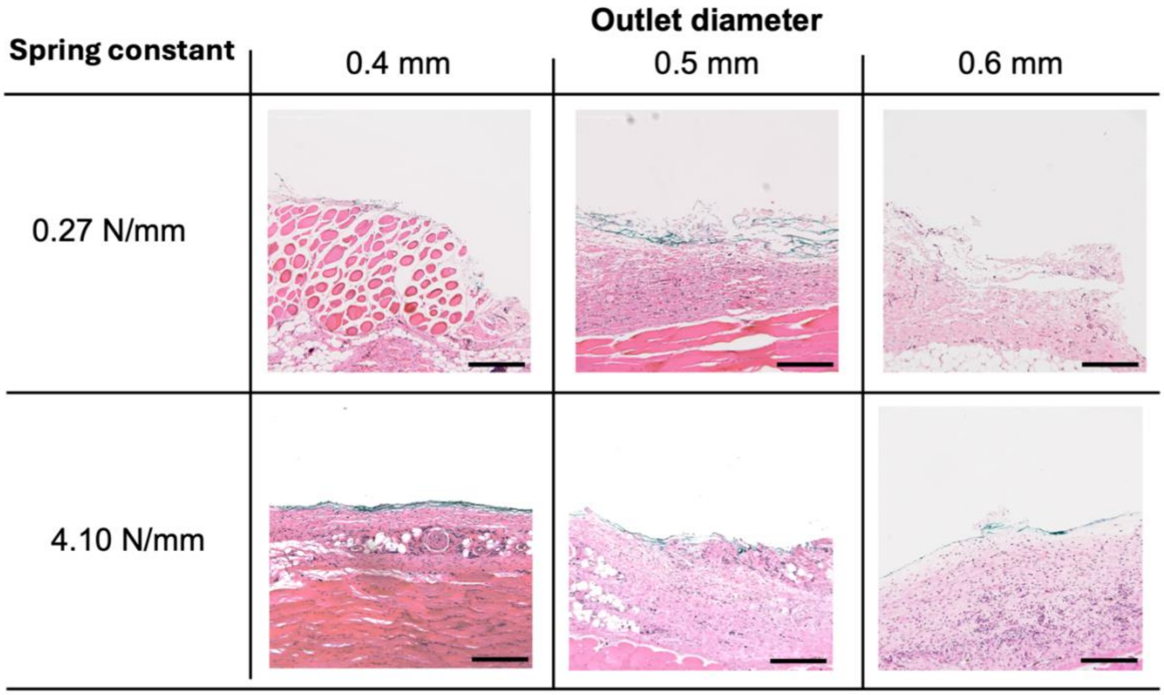


**Fig. S2. Histological safety examination of jet injection on ex vivo rat skin.** Hematoxylin and eosin (H&E) slides from the weakest and strongest jetting devices loaded with green tissue-marking dye and actuated onto ex vivo rat skin. The dye is deposited within the hypodermis without penetrating the dermis or epidermis (scale bar, 300 μm).

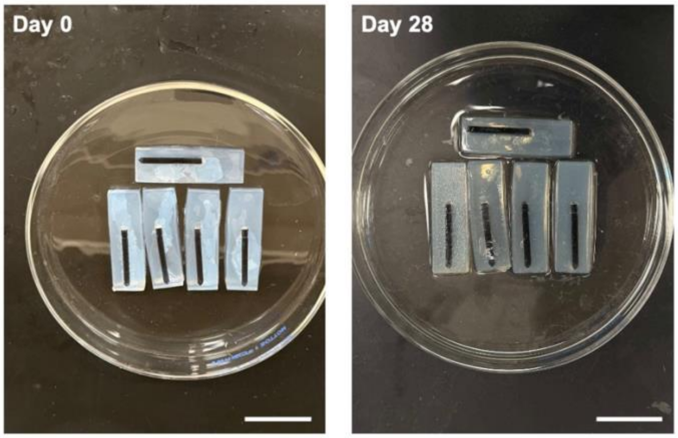


**Fig. S3. Actuator stability.** Devices embedded in a tissue-mimicking hydrogel incubating at 37°C and 50 RPM show no premature actuation for 28 days. The black coating is a ceramic coating to prevent moisture ingress. (n=5 devices) (scale bar, 3 cm).

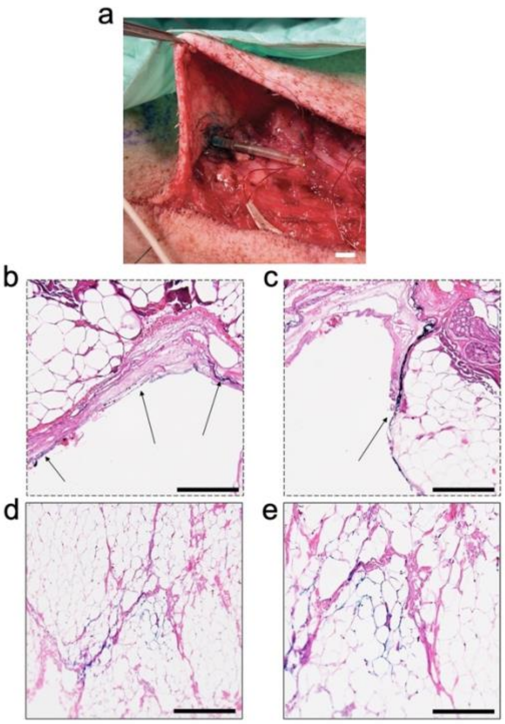


**Fig. S4. Jet injection and histological safety examination in swine. a,** RES-Q implant placement and actuation in swine (scale bar, 6 mm). **b-e,** Representative histological images of dye deposition following RES-Q implant actuation in swine. The dye is deposited within the hypodermis without penetrating the dermis or epidermis (b,c,e scale bar, 200 μm; d scale bar, 300 μm).

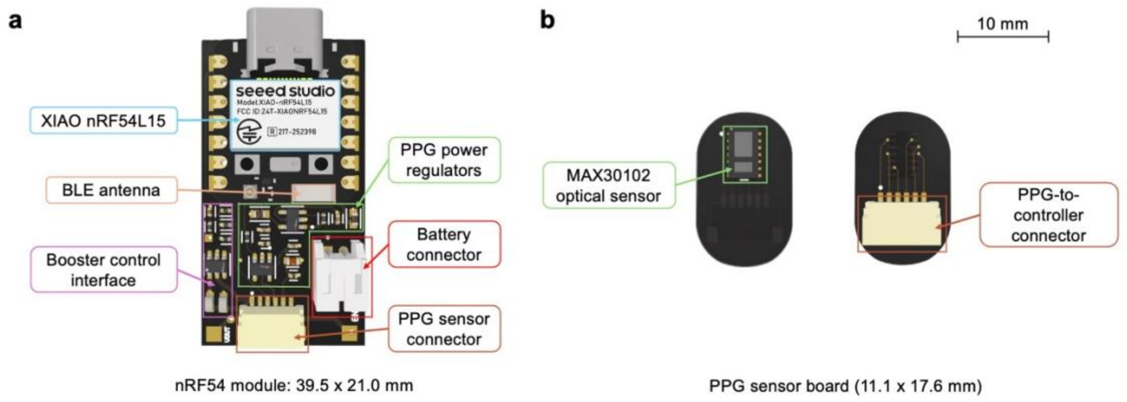


**Fig. S5. Printed circuit board layout of the wearable sensing and voltage booster system. a,** The main board with the nRF54 module integrates the photoplethysmography (PPG) regulator, voltage actuation circuit, microcontroller unit (MCU), Bluetooth Low Energy (BLE) antenna, and battery connector, along with the voltage actuation circuit and voltage booster. **b,** Separate MAX30102 optical sensor board used for rat-paw photoplethysmography and connected to the main board.

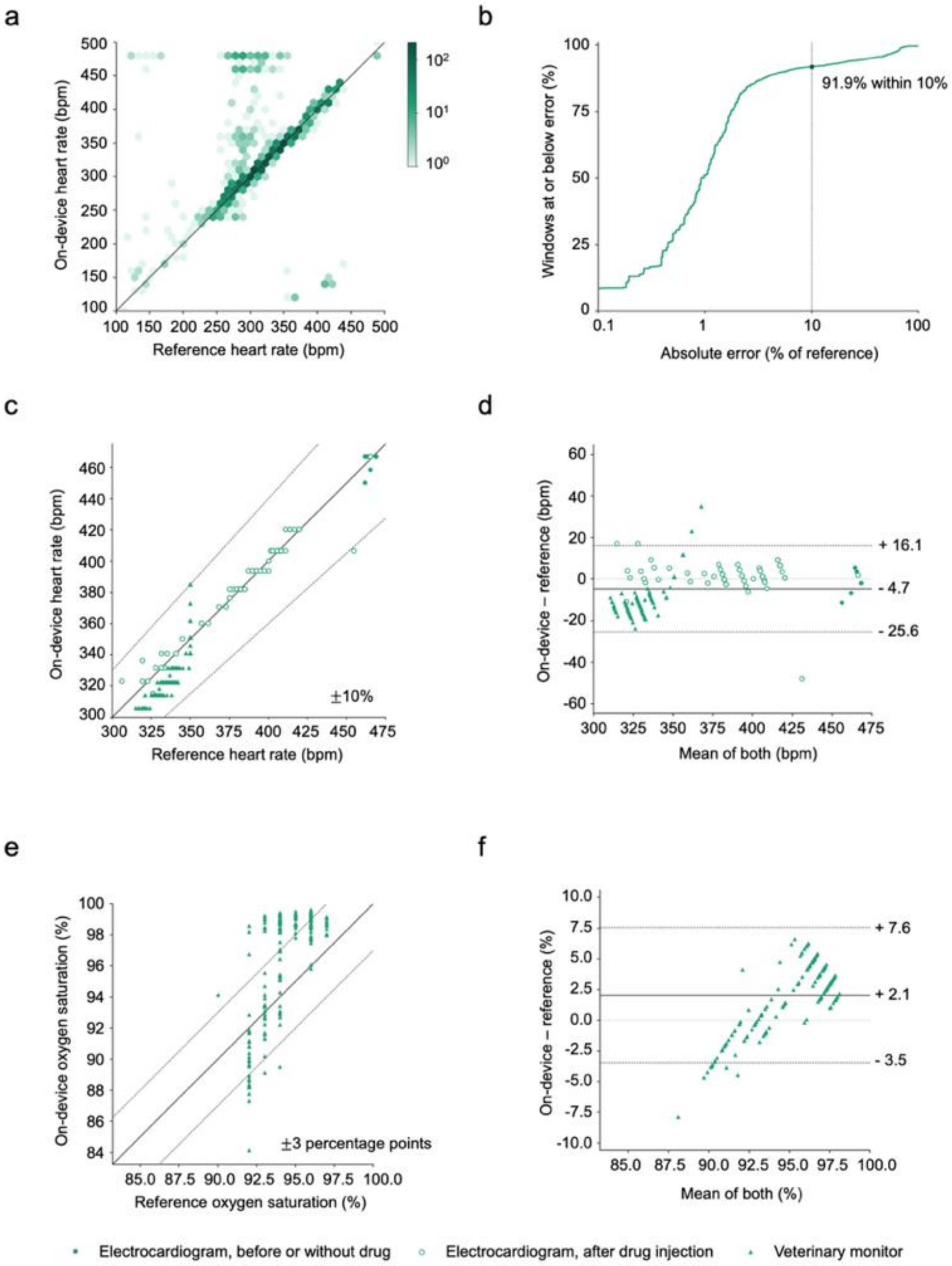


**Fig. S6. Validation of on-device heart-rate estimation against reference analyses and electrocardiography.** Validation of on-device heart-rate estimation against reference analyses and electrocardiography. **a,** Comparison of heart-rate estimates computed on the wearable with an offline spectral reference across held-out recordings

(2,468 10-s windows not used for algorithm tuning). Color intensity indicates the number of windows, and the solid line denotes identity. **b,** Cumulative distribution of absolute error of the same estimates, as a percentage of the reference. Half of the windows fall within 1% (3 beats/min at the median reference rate of 321 beats/min) and 91.9% within 10%. **c,** Comparison of photoplethysmography (PPG) derived on-device heart rate with simultaneously recorded electrocardiogram-derived heart rate (circles) and a veterinary monitor (triangles). Dashed lines indicate $\pm$10% of the reference. Measurements comprise 245 windows from 5 rats (one rat with a simultaneously recorded electrocardiogram and four rats with the veterinary monitor). **d,** Bland-Altman analysis of the same heart rate measurements. The mean difference between on-device and reference heart rate was -7.1 beats/min, with limits of agreement from -24.6 to +10.3 beats/min. **e,** Comparison of PPG-derived on-device oxygen saturation with the veterinary patient monitor. Dashed lines indicate $\pm$3% percentage points of the monitor reference. Measurements comprise 168 windows from 4 rats with the veterinary monitor. **f,** Bland-Altman analysis of the same oxygen saturation measurement. The mean difference between on-device and monitor oxygen saturation was +2.1%, with limits of agreement from -3.5 to +7.6%.

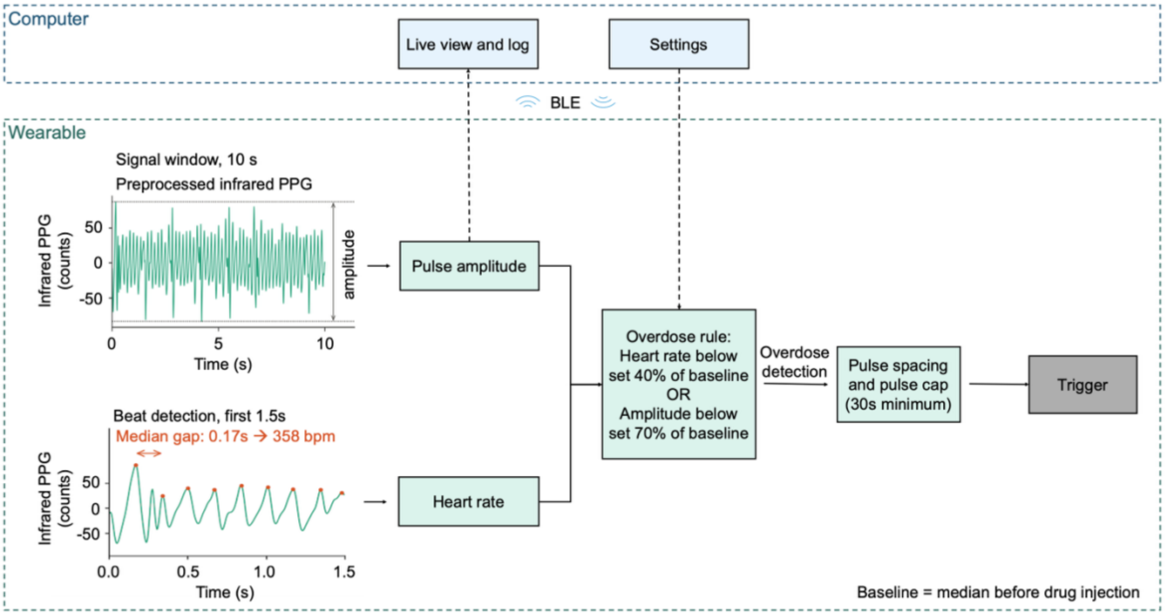


**Fig. S7. Overdose detection algorithm and triggering workflow.** Block diagram of on-device closed-loop workflow for photoplethysmography (PPG) processing, overdose detection, and trigger control. Bluetooth Low Energy = BLE.

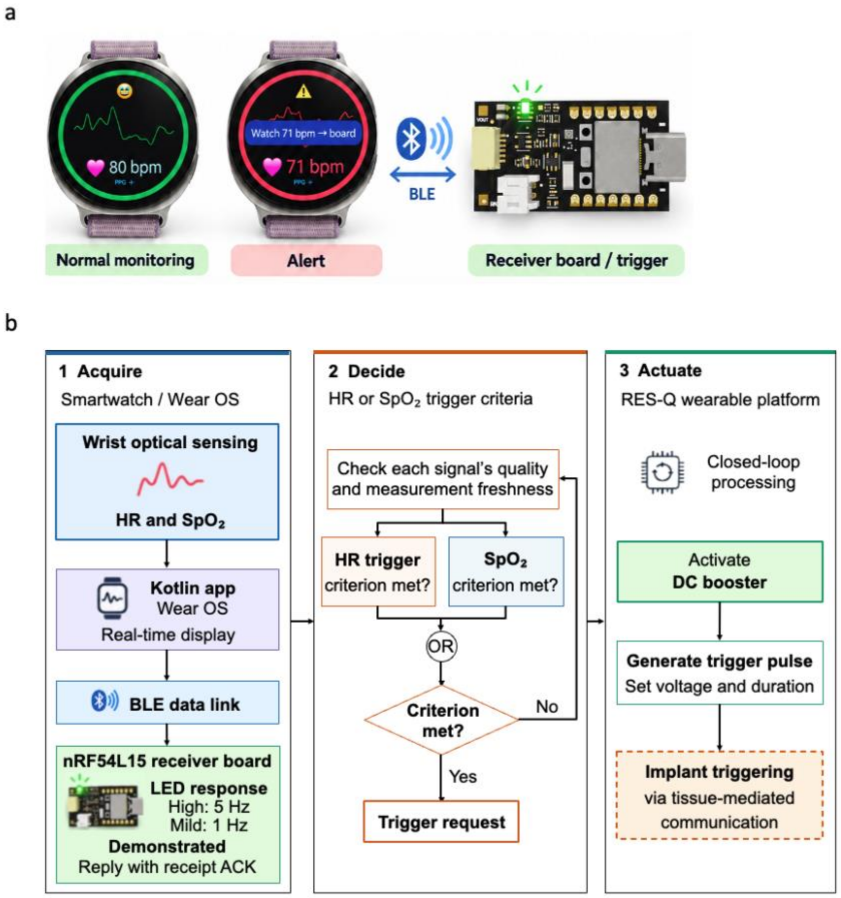


**Fig. S8. RES-Q platform integration with a smartwatch. a,** Smartwatch interface during normal monitoring and alert states for the receiver. **b,** Data pathway from smartwatch optical sensing and the Wear OS application through Bluetooth Low Energy to the RES-Q wearable platform for downstream processing and triggering.

a

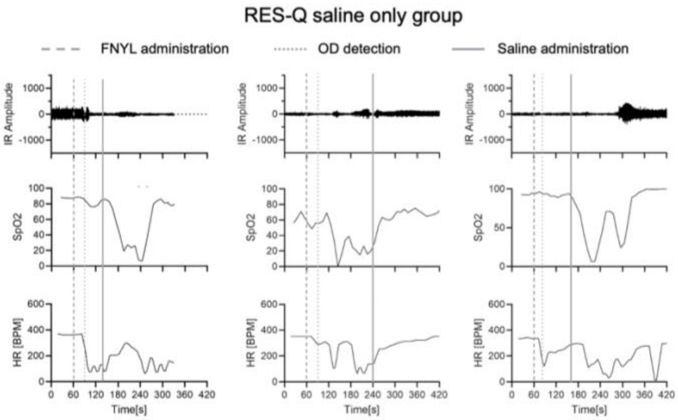
RES-Q saline only group
FNYL administration
OD detection
Saline administration
IR Amplitude
SpO2
HR [BPM]
Time[s]

b

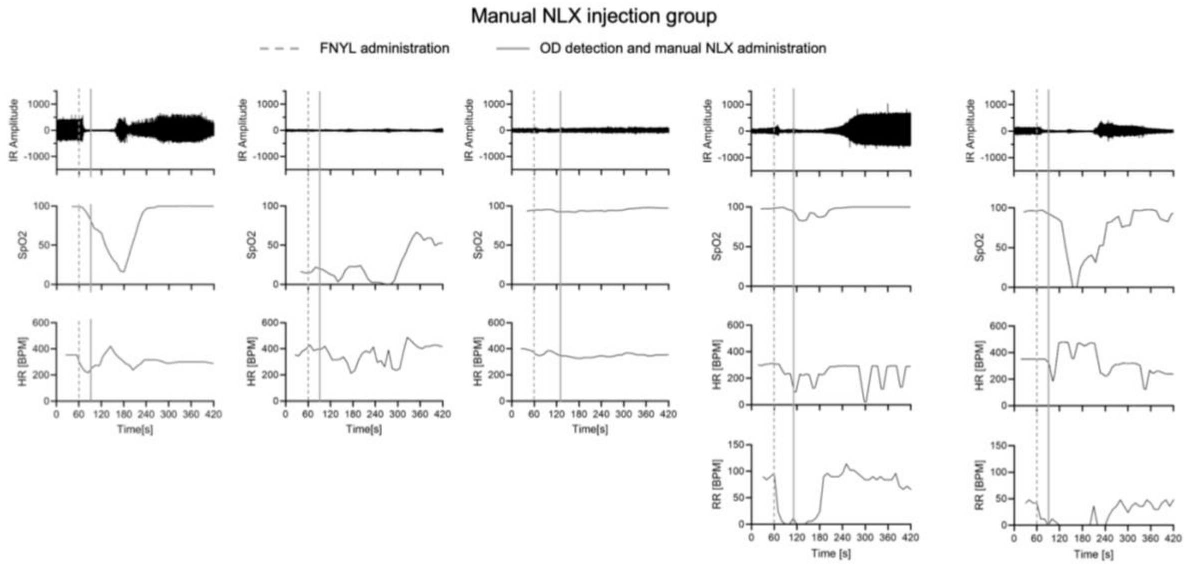
Manual NLX injection group
FNYL administration
OD detection and manual NLX administration
IR Amplitude
SpO2
HR [BPM]
RR [BPM]
Time[s]
Time [s]

c

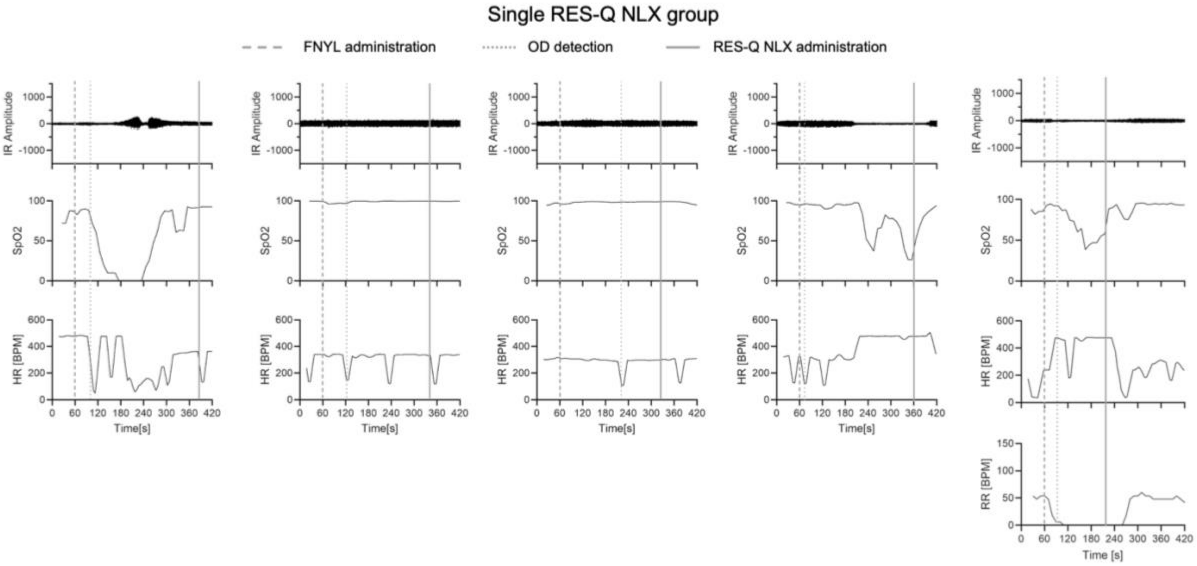
Single RES-Q NLX group
FNYL administration
OD detection
RES-Q NLX administration
IR Amplitude
SpO2
HR [BPM]
RR [BPM]
Time[s]
Time [s]

**d**

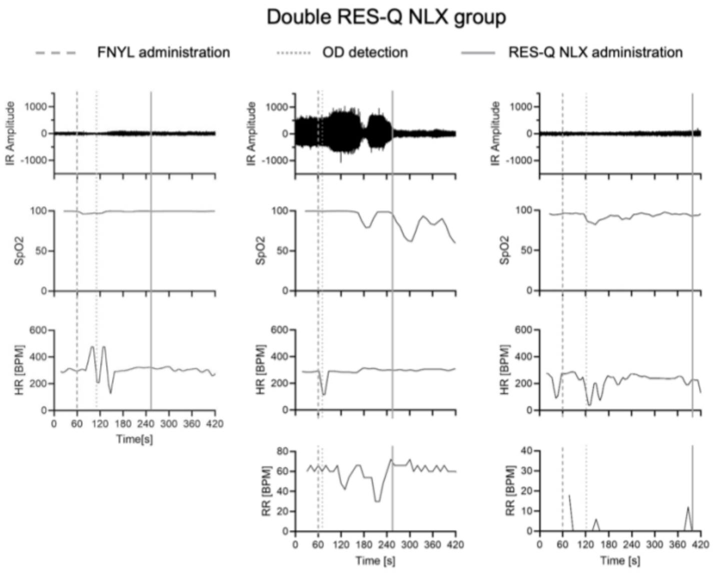


**Fig. S9. Fentanyl opioid overdose rescue data.** Vital signs collected during a fentanyl (FNYL) induced overdose (OD) for: **a,** a negative control group where rats were deployed with a RES-Q device loaded with saline only (*n* = 3 rats). **b,** a positive control where rats received a manual injection of naloxone (NLX) (*n* = 5 rats). **c,** a group where rats were deployed with a single RES-Q device loaded with naloxone. Vital signs in the last graph can also be found in Figure 4b and is presented again here for context. (*n* = 5 rats). **d,** a group where rats were deployed with two RES-Q devices loaded with naloxone (NLX) to demonstrate sequential dosing capabilities (*n* = 3 rats). Each column represents a different animal trial. Heart Rate = HR. Infrared = IR. Peripheral capillary oxygen saturation = $SpO_2$.

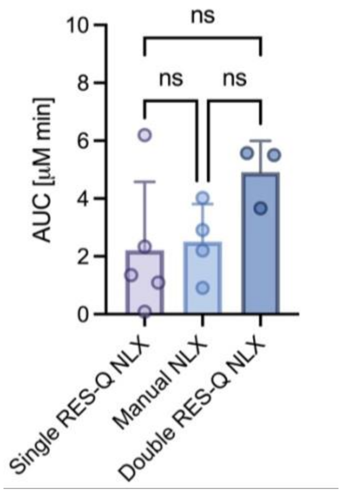


**Fig. S10. Pharmacokinetic area under the curve (AUC) for the groups receiving naloxone.** AUC comparison between animals receiving a single dose of naloxone (NLX) from the RES-Q device (*n* = 5 rats), animals receiving a manual injection of naloxone (*n* = 4 rats), and animals that received sequential doses of naloxone from the RES-Q network (*n* = 3 rats). Data represent means ± s.d and individual data. Statistical significance of difference between groups was computed via ordinary one-way analysis of variance (ANOVA), $2.263_{(2,9)} = 78.04$ (P =0.1599). ns = no statistically significant difference.

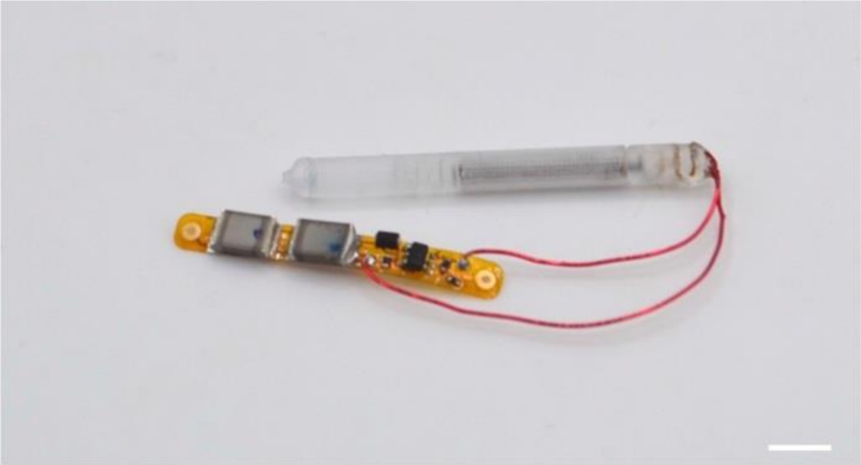

**Fig. S11. RES-Q device loaded with liquid epinephrine.** A clinically relevant dose of 0.3 mg of epinephrine in 16 µL of saline is loaded into the implant, highlighting a potential use case for the treatment of severe allergic reactions (scale bar, 4 mm).

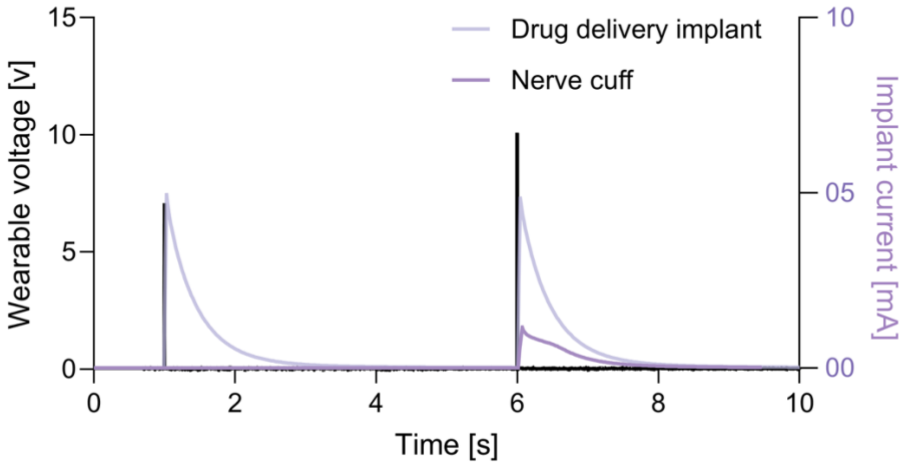


**Fig. S12. Sequential in-body communication triggering of the RES-Q implant and the Smart Wireless Autonomous Networking System (SWANS) neurostimulator implant.** Demonstration of potential for coordinated, targeted pharmacological and electrical interventions, with devices actuated in ex vivo chicken breast tissue.

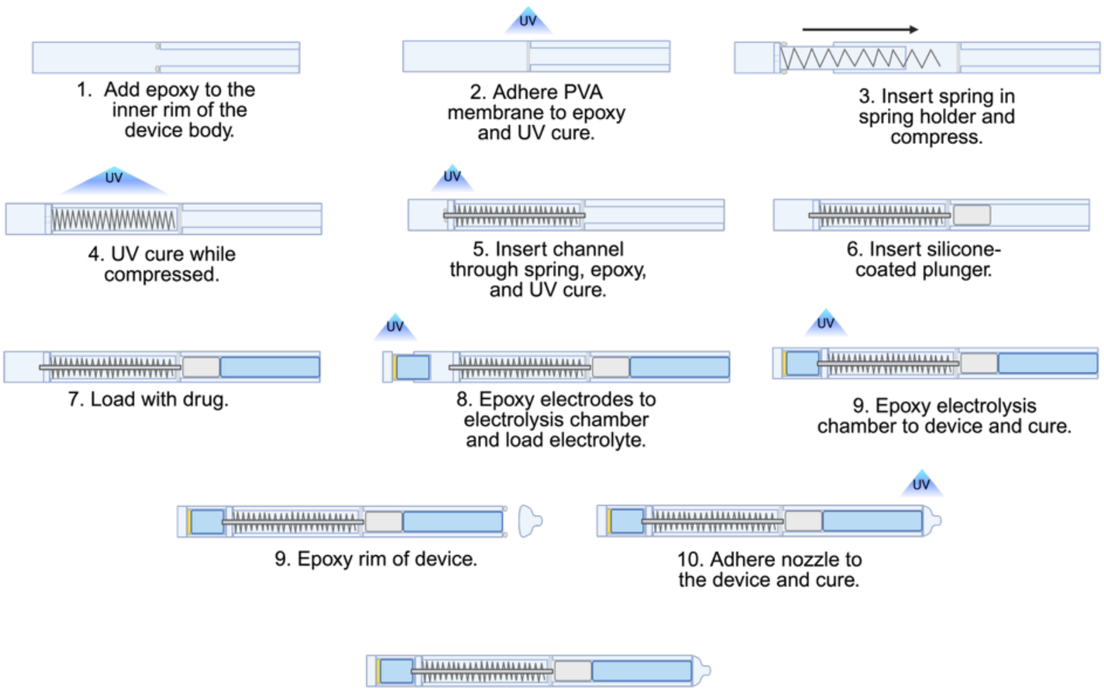


**Fig. S13. Device assembly.** Step-by-step process for assembling the drug delivery device. Illustrations created with BioRender.com. Polyvinyl alcohol = PVA. Ultraviolet = UV.

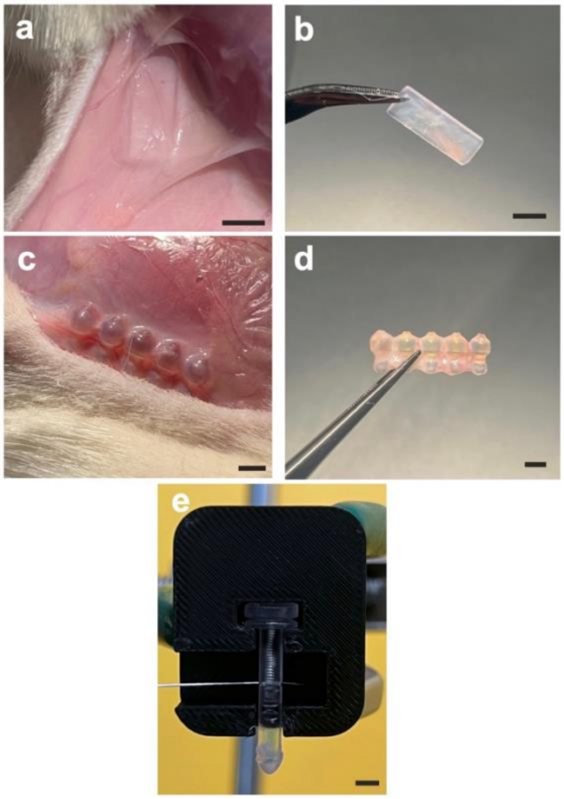


**Fig. S14. Devices for fibrosis harvesting. a-b**, Subcutaneous rectangle implant possessing fibrotic tissue that was harvested to use during tensile tests (scale bar, 5 mm). **a,** Implant in the subcutaneous dorsum of a Sprague Dawley rat. **b,** Implant following explanation. **c-d**, Subcutaneous nozzle implant possessing fibrotic tissue that was harvested to use during ex vivo jetting studies (scale bar, 3 mm). **c,** Implant in the subcutaneous dorsum of a rat. **d,** Implant following explanation. **e**, Fibrotic tissue attached to jetting apparatus during *ex vivo* study (scale bar, 3 mm).

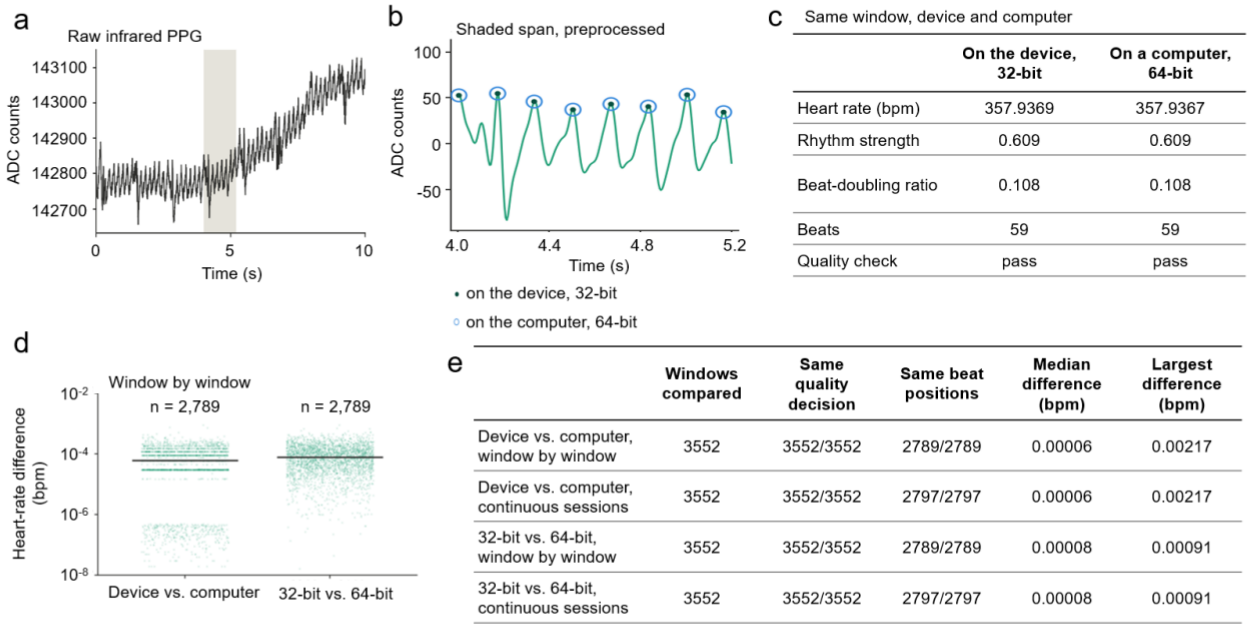


| | On the device, 32-bit | On a computer, 64-bit |
|---|---|---|
| Heart rate (bpm) | 357.9369 | 357.9367 |
| Rhythm strength | 0.609 | 0.609 |
| Beat-doubling ratio | 0.108 | 0.108 |
| Beats | 59 | 59 |
| Quality check | pass | pass |

| | Windows compared | Same quality decision | Same beat positions | Median difference (bpm) | Largest difference (bpm) |
|---|---|---|---|---|---|
| Device vs. computer, window by window | 3552 | 3552/3552 | 2789/2789 | 0.00006 | 0.00217 |
| Device vs. computer, continuous sessions | 3552 | 3552/3552 | 2797/2797 | 0.00006 | 0.00217 |
| 32-bit vs. 64-bit, window by window | 3552 | 3552/3552 | 2789/2789 | 0.00008 | 0.00091 |
| 32-bit vs. 64-bit, continuous sessions | 3552 | 3552/3552 | 2797/2797 | 0.00008 | 0.00091 |

**Fig S15. Agreement between on-device and reference photoplethysmography (PPG) processing. a**, Representative 10-s infrared PPG window processed by the on-device algorithm. The shaded region is enlarged in b. **b**, Preprocessed waveform showing beat locations identified by the 32-bit on-device and 64-bit reference implementations. **c**, Comparison of heart-rate and signal-quality outputs for the representative window. **d,** Heart-rate differences between implementations across individual windows. **e**, Agreement in heart-rate estimates, beat locations, and quality decisions across window-wise and continuous-session processing. Beat locations and quality decisions were identical in all comparable windows, with negligible differences in heart-rate estimates. Beats per minute = BPM.

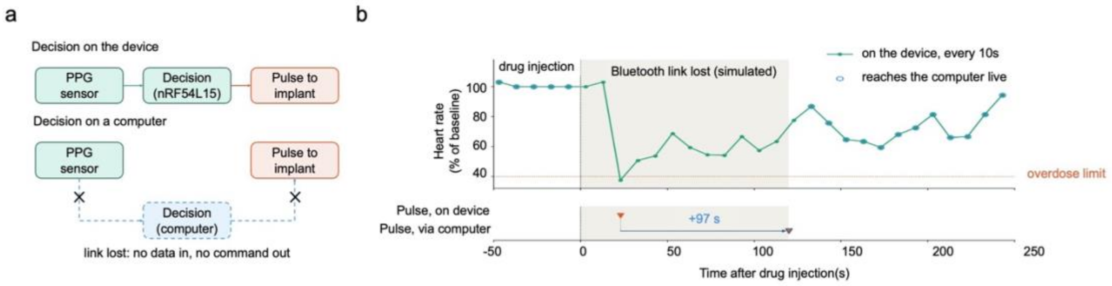


**Fig. S16. Robustness of autonomous triggering to Bluetooth link interruption.**

**a**, Schematic comparison of on-device and computer-dependent overdose-detection architectures. When the decision is executed locally on the nRF54L15, loss of the BLE link does not interrupt implant triggering. In contrast, when the decision depends on an external computer, BLE interruption prevents receipt of live physiological data and delays pulse delivery. **b**, Representative replay of an overdose session with simulated BLE link loss using a heart rate baseline drop to 40% of baseline as the overdose detection threshold. Heart rate (% of baseline) is evaluated locally every 10 s on the device (green), while values reaching the computer are shown as blue circles when communication is available. The on-device controller detects the overdose condition and delivers the pulse during the link-loss interval, whereas a computer-dependent pathway would deliver the pulse only after communication recovery, causing a 97-s delay. Photoplethysmography = PPG.

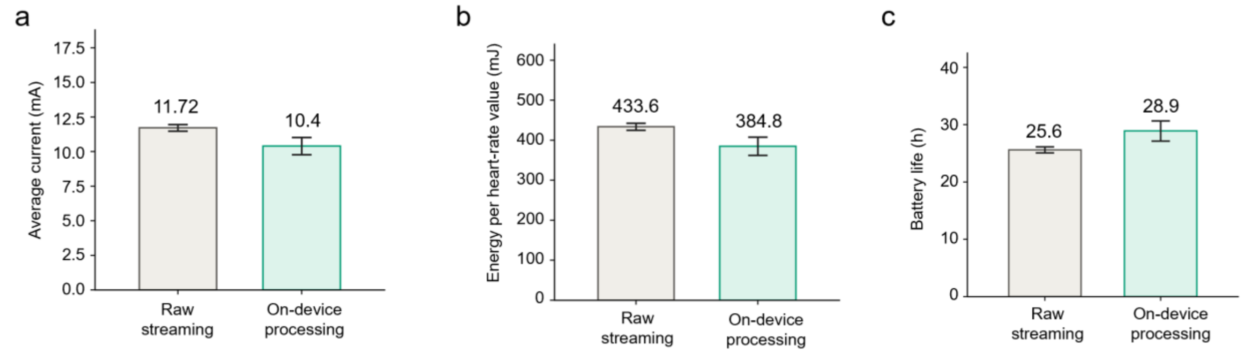


**Fig. S17. Estimated wearable power consumption and nominal runtime in two operating modes.** Both modes were evaluated using the same 300-mAh lithium-polymer cell and the same charge-termination criterion. **a**, Estimated session-average current during raw streaming and on-device processing. Current was estimated from the change in battery state of charge over each session. **b**, Estimated energy consumed per 10-s analysis window, calculated as the session-average current multiplied by the nominal cell voltage of 3.7 V and the 10-s analysis interval. **c**, Estimated continuous runtime per charge for a nominal 300-mAh cell, calculated as the nominal cell capacity divided by the session-average current. One session was recorded per mode. *n* = 5 sessions per mode.
